# KDM2B is required for ribosome biogenesis and its depletion unequally affects mRNA translation

**DOI:** 10.1101/2024.05.22.595403

**Authors:** Vollter Anastas, Evangelia Chavdoula, Alessandro La Ferlita, Burak Soysal, Ilaria Cosentini, Giovanni Nigita, Michael G. Kearse, Philip N. Tsichlis

**Author notes:** Correspondence: Philip N. Tsichlis, Vollter Anastas. **Conflict of Interest Statement:** PNT is a co-founder of “Epi-Cure” which specializes in demethylase inhibitors. The other authors have no conflicts of interest.

## Abstract

KDM2B is a JmjC domain lysine demethylase, which promotes cell immortalization, stem cell self-renewal and tumorigenesis. Here we employed a multi-omics strategy to address its role in ribosome biogenesis and mRNA translation. These processes are required to sustain cell proliferation, an important cancer hallmark. Contrary to earlier observations, KDM2B promotes ribosome biogenesis by stimulating the transcription of genes encoding ribosome biogenesis factors and ribosomal proteins, particularly those involved in the biogenesis of the 40S ribosomal subunits. Knockdown of KDM2B impaired the assembly of the small and large subunit processomes, as evidenced by specific defects in pre-ribosomal RNA processing. The final outcome was a decrease in the rate of ribosome assembly and in the abundance of ribosomes, and inhibition of mRNA translation. The inhibition of translation was distributed unequally among mRNAs with different features, suggesting that mRNA-embedded properties influence how mRNAs interpret ribosome abundance. This study identified a novel mechanism contributing to the regulation of translation and provided evidence for a rich biology elicited by a pathway that depends on KDM2B, and perhaps other regulators of translation.

## INTRODUCTION

*KDM2B* encodes a jumonji C (JmjC) domain-containing lysine demethylase, which targets histone H3K36me2/me1 and perhaps histone H3K4me3 and histone H3K79 me3/me2 and functions as an oncogene in several types of tumors. Following the original demonstration of its oncogenic potential in Moloney murine leukemia virus (MoMuLV)-induced rodent T cell lymphomas (1), KDM2B was shown to also promote the development of human lymphoid and myeloid malignancies(2, 3), as well as bladder cancer (4), pancreatic cancer (5), breast cancer (6), gliomas (7), and prostate cancer (8). In breast cancer, its highest expression was observed in the basal-like triple negative variant (TNBC), an aggressive variant, which is more common in younger women and comprises about 15-20% of all breast cancers (9). In these tumors, KDM2B represses the expression of a set of microRNAs which target multiple components of the polycomb repressive complexes PRC1 and PRC2, promoting cancer stem cell self-renewal (6). Additionally, KDM2B protects cells from oxidative stress (10), inhibits senescence (1, 11), promotes cell proliferation and invasiveness (1, 4, 11), and inhibits apoptosis (4, 6). It also promotes angiogenesis (4), and our more recent studies showed that it is a master regulator of intermediary metabolism (12). In this report, we present evidence that KDM2B has an important role in the regulation of ribosomal biogenesis and mRNA translation.

Ribosomal biogenesis is a well-orchestrated multistep process, which is initiated with the synthesis of 47S pre-ribosomal RNA (pre-rRNA) from a set of approximately 400 head to tail tandem ribosomal DNA repeats (13). These repeats, also known as nucleolar organizer regions (NORs), map in the short arms of the acrocentric chromosomes 13, 14, 15, 21 and 22 (13). Pre-rRNA transcribed from these repeats, recruits a large number of proteins and noncoding RNAs, triggering the self-assembly of the nucleolus (14). The first stable structure in the biogenesis of ribosomal subunits, is the 90S pre-ribosome, a very large structure, which ultimately gives rise to both the 40S and the 60S subunits (15). The maturation process depends on the ordered cleavage of the 47S pre-rRNA, at multiple sites, which is regulated by precisely orchestrated, protein-guided RNA and protein movements. These cleavage events, combined with several 5′ to 3′ and 3′ to 5′ exonuclease digestions they are coupled to, remove two external transcribed spacers (5′ and 3′ ETS) and two internal transcribed spacers (ITS1 and ITS2) to give rise to the mature 18S, 5.8S and 28S ribosomal RNAs (16). Critical in this process is the role of the 5′ ETS, which in mammalian cells is approximately 3.5 kb long, and which is removed by 3 consecutive cleavage events, the A′ cleavage, which occurs co-transcriptionally (17), and the A0 and A1 cleavages, which result in the stepwise removal of the entire 5′ ETS (16, 18). The separation of the small subunit processome (SSU) from the precursor 60S particle occurs after cleavage 2 in ITS1.

Perturbations in ribosomal biogenesis may alter the number and the composition of the ribosomes, impacting the rate and fidelity of translation (19, 20). However, given that the cellular transcriptome is composed of mRNAs with diverse features, such changes may influence unequally the translation of different mRNAs (21). Translation initiation of most mRNAs depends on the m7G cap at their 5′ end. Capped mRNAs are also heterogeneous, some of them having longer coding sequences (CDS) than others, with the ones with longer CDSs tending to have long highly structured 5′ untranslated regions (5′ UTRs) (22). Additionally, some mRNAs have short open reading frames in their 5′UTRs (23), or binding sites for select RNA binding proteins at their 5′ or 3′ UTRs (24). Differential reading of these features in an environment of changing ribosome numbers or changing ribosome composition could affect translation of mRNAs unequally (21, 25).

Earlier studies had suggested that KDM2B encodes a nucleolar protein which inhibits ribosomal biogenesis by repressing the transcription of pre-ribosomal RNA (26). This finding was unexpected because KDM2B generally promotes cell proliferation, which requires enhancement rather than repression of ribosomal biogenesis. In this report, we revisited this important question with unbiased multi-omics studies in control and shKDM2B-transduced MDA-MB-231 cells, and the results showed that KDM2B strongly promotes ribosomal biogenesis and mRNA translation, as expected. Specifically, KDM2B promotes transcriptionally the expression of many ribosomal proteins and ribosome biogenesis factors (RBFs) with stronger effects on the expression of factors associated with the small subunit processome (SSU) and the assembly of the 40S ribosomal subunit. Consistent with these findings, the knockdown of KDM2B impairs primarily the efficiency of the A0 cleavage in the 5′ ETS of the 30S pre-ribosomal RNA, which occurs during the transition of the SSU processome from the pre-A1 to the post-A1 stage (27). Another cleavage, which was weakly impaired in the shKDM2B cells, is cleavage 4 in ITS2. Defects in the processing of pre-ribosomal RNA and in ribosome biogenesis, are primarily due to changes in the expression of ribosomal protein and RBF-encoding genes, which are regulated by KDM2B via ncPRC1.1-dependent mechanisms, and in concert with MYC. ncPRC1.1 is a variant of Polycomb Complex 1 (PRC1), and KDM2B is the DNA/chromatin-targeting component of the complex. Inhibition of the preceding events by KDM2B knockdown, decreases the rate of ribosome assembly and maturation and the overall abundance of ribosomes, and results in inhibition of global mRNA translation. Most important however, the inhibition of mRNA translation is selective, with significant variation in the efficiency of translation of mRNAs with distinct structural features. These data support the hypothesis that changing ribosome numbers may be interpreted differently by mRNAs. The differential interpretation of these changes by mRNAs with different structural features provides the basis for an unexpectedly rich biology, which may be elicited by perturbations in the expression and/or function of KDM2B and other regulatory factors.

## RESULTS

### KDM2B is overexpressed in basal-like breast cancer, and its knockdown impacts the expression of genes involved in ribosome biogenesis and translation

*KDM2B* is expressed at higher levels in breast cancer than in the normal mammary gland, but its highest expression was observed in the basal-like subtype (Fig. S1A and S1B). In agreement with this observation, the difference in KDM2B expression between breast cancer and the adjacent normal tissue was more robust in basal like breast cancer (Fig. S1C). Additionally paired comparisons of KDM2B expression in tumor tissue and the adjacent normal mammary gland, revealed that the higher expression of KDM2B in the tumor tissue was significant in basal like, than in other breast cancer subtypes suggesting the increased level is more specific to basal-like breast cancer (Fig S1D).

To determine the functional role of KDM2B in basal like breast cancer, we compared the transcriptomes of control and KDM2B knockdown MDA-MB-231 cells (12), using Gene Set Enrichment Analysis (GSEA) (28), and focusing on the Gene Ontology (GO) domain “Biological Processes”. This analysis revealed significant negative enrichment in “ribosome biogenesis”, “cytoplasmic translation”, “rRNA metabolic process”, as well as in “ncRNA processing” and “ribosomal large subunit biogenesis” in KDM2B knockdown cells (Figure 1A and 1B). Given that cancer cells are highly dependent on mRNA translation (19), this observation is consistent with the role of KDM2B in oncogenesis. The heatmap in figure 1C, shows the differential expression of genes encoding ribosomal proteins and ribosome biogenesis factors (RBFs) (GO ribosomal biogenesis), in control and shKDM2B-transduced cells. Genes shown in the heatmap include those whose differential expression in control and shKDM2B cells meets an adjusted p value of <0.05 and a threshold of absolute Fold Change (Control/shKDM2B) of log_2_ ≥ 0.58.

**Figure 1.**
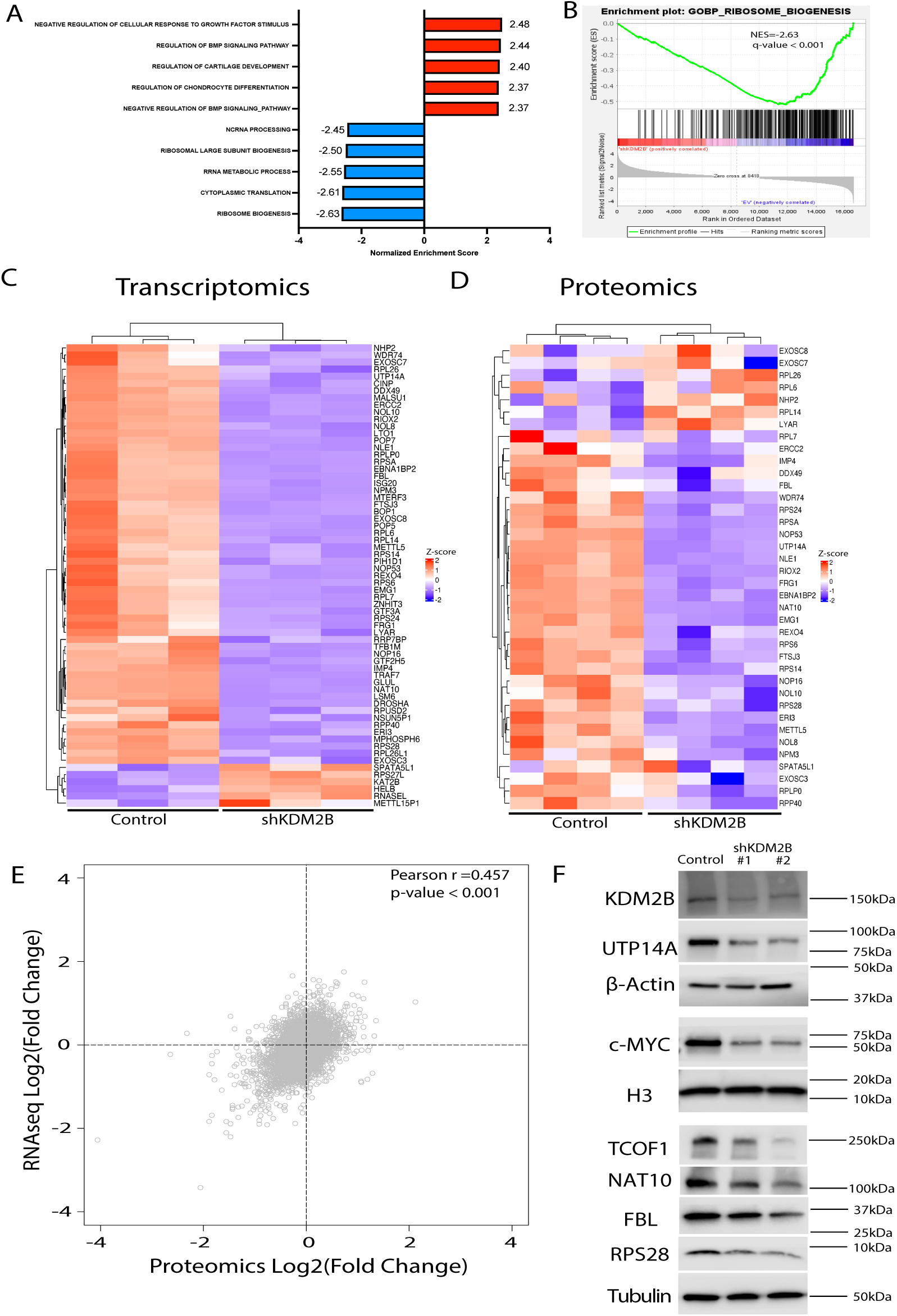
Knockdown of KDM2B downregulates genes involved in ribosomal biogenesis and mRNA translation. **A**. GSEA analysis showing the normalized enrichment scores of the top 5 positively enriched and the top 5 negatively enriched gene ontology terms in KDM2B knockdown MDA-MB-231 cells. **B.** Gene set enrichment plot of Ribosome Biogenesis. **C.** Heatmap showing the relative RNA levels of all ribosome biogenesis genes whose expression was altered (Log2FC<=+/-0.58, FDR<0.05), in shKDM2B MDA-MB-231 cells. **D**. Heatmap showing the relative abundance of proteins involved in ribosome biogenesis whose expression was altered in shKDM2B MDA-MB-231 cells. Protein abundance was determined by TMT proteomics analysis. Unbiased clustering in C and D was performed with the Complex Heatmap R package (https://github.com/jokergoo/ComplexHeatmap). Normalized signal intensities for all genes were based on Z-scores, calculated from the geometric mean of the expression of each gene. **E.** Scatter plot comparing the log2(fold change) of the expression of individual genes at the RNA and protein levels in shKDM2B relative to control cells. Pearson correlation coefficient 0.457, p-value < 0.001. **F.** Western blots comparing the protein levels of selected ribosome biogenesis factors in control and shKDM2B MDA-MB-231 cells.

To determine whether differences in the abundance of gene transcripts reflect differences in protein abundance, we carried out quantitative Tandem Mass Tag (TMT) proteomics, using lysates of the same cells, and the results revealed significant agreement with the transcriptomic data (Fig 1D), consistent with the partial concordance between the transcriptomic and proteomic data (Fig 1E). The expression of a subset of genes encoding ribosomal proteins (RPS28), or factors involved in the transcription and processing of ribosomal RNA (MYC, UTP14A, TCOF1, NAT10 and FBL), was also examined by western blotting and the results were consistent with the transcriptomic and proteomic data (Fig 1F).

### KDM2B regulates the transcription of ribosome biogenesis genes by functioning in the context of ncPRC1.1 and in concert with MYC

The decreased steady state levels mRNAs encoding ribosomal proteins and RBFs in shKDM2B cells (Fig 1A-C) suggested that these may be regulated by KDM2B at the level of transcription. The exon intron split analysis (29) of our RNA-Seq data supported this conclusion by showing that the abundance of intron reads of genes encoding RBFs correlates well with the abundance of exon reads (Fig 2A). KDM2B is a lysine-specific demethylase which has been linked to both transcriptional activation and transcriptional repression (5, 30, 31). Transcriptional repression is mediated primarily by EZH2, the catalytic component of PRC2, which is positively regulated by KDM2B and promotes trimethylation of histone H3 at K27 (6, 32). Transcriptional activation on the other hand, has been linked to the ncPRC1.1 complex, which is targeted to DNA by KDM2B, one of its components, and indirectly promotes the acetylation of histone H3 at K27 (31, 33, 34). Our earlier studies had shown that the KDM2B-dependent positive regulation of genes encoding multiple metabolic enzymes is independent of EZH2, but depends on ncPRC1.1, whose integrity is compromised in shKDM2B cells because of the downregulation of several of its members, including KDM2B itself, USP7, TRIM27, SKP1 and RYBP (12). Here we show that the regulation of ribosome biogenesis genes by KDM2B is also EZH2 independent (Fig 2B). In parallel experiments we compared the expression of the same RBFs in shPCGF1 or shPCGF4/BMI1-transduced MDA-MB-231 cells. PCGF1 is a component of ncPRC1.1, while PCGF4/BMI1 is a component of the canonical PRC1. The results showed that whereas the targeting of ncPRC1.1 (shPCGF1) phenocopies the KDM2B knockdown phenotype, the targeting of the canonical PRC1 (shPCGF4) does not (Fig 2C). The fact that the depletion of PCGF1, which like KDM2B is a member of the ncPRC1.1 complex, phenocopies the effects of the depletion of KDM2B on the expression of the set of genes in figure 2C, strongly suggests that KDM2B regulates the expression of these genes, by functioning in the context of ncPRC1.1.

**Figure 2.**
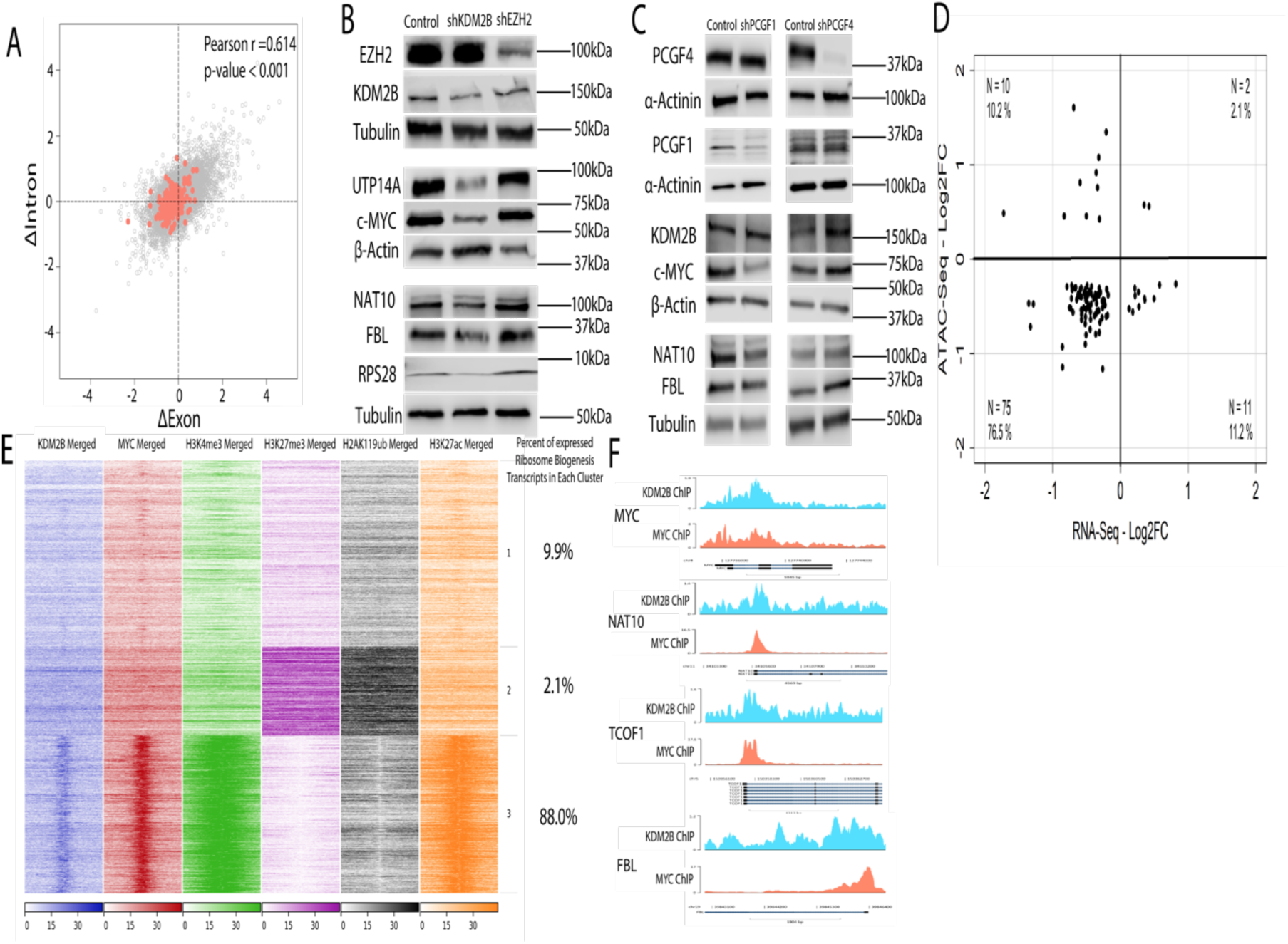
KDM2B regulates ribosome biogenesis genes transcriptionally by functioning in the context of ncPRC1.1 and in concert with MYC. **A.** Exon-Intron split analysis of the RNA-Seq data in control and shKDM2B MDA-MB-231 cells. The scatter plot compares the log2(fold change) of exon and intron reads of individual genes in shKDM2B relative to the control cells. Orange dots correspond to ribosome biogenesis genes. Pearson correlation coefficient, 0.614, p-value < 0.001. **B and C.** Western blot analysis of the proteins encoded by a set of KDM2B-dependent ribosome biogenesis genes in control and shEZH2, shPCGF1, and shPCGF4-transduced MDA-MB-231 cells. **D.** Scatter plot comparing the log2(fold change) in RNA levels with the log2(fold change) in chromatin accessibility. in the transcription start site (TSS) of ribosome biogenesis genes. Chromatin accessibility was measured by ATAC-Seq. **E.** FLUFF clustering of the TSS-centered mapped reads generated by the indicated ChIP-Seq experiments in MDA-MB-231 cells produced three distinct clusters. Percentages of the most abundant isoform of expressed of ribosomal biogenesis factors based on RNA-seq data. **F.** Alignment of the KDM2B and MYC ChIPseq peaks in the promoters of MYC, NAT10, TCOF1, and FBL.

The assay for transposase-accessible chromatin with sequencing (ATAC-Seq) in control and shKDM2B-transduced MDA-MB-231 cells, was used to identify KDM2B/ncPRC1.1-induced changes in the chromatin accessibility of ribosome biogenesis genes. Our earlier studies, focusing primarily on the effects of shKDM2B on the accessibility of genes encoding metabolic enzymes and their regulators like MYC and ATF4, had shown that the knockdown of KDM2B results in decreased chromatin accessibility in 25,312 sites, and increased accessibility in 12,037 sites (absolute Log2FC ≥ 0.58, FDR < 0.05), and that changes in chromatin accessibility correlate with changes in gene expression (12). Here we focused on the chromatin accessibility of ribosome biogenesis genes and the results confirmed the overall concordance between decreasing accessibility and decreasing gene expression in shKDM2B cells (Fig 2D).

FLUFF analysis (35) of all the mapped reads in our ChIP-Seq study of KDM2B, H3K4me3, H2AK119ub, H3K27me3, H3K27ac, and MYC in MDA-MB-231 cells (12) showed that cellular genes fall into three distinct clusters. One of these clusters contains actively transcribed genes, which are characterized by high abundance of H3K4me3 and H3K27ac marks, and by the concerted binding of KDM2B and MYC in the area flanking the Transcription start site (TSS). The other two clusters were characterized by the low binding of KDM2B and MYC and either by the high abundance of chromatin marks associated with transcriptional repression, or by the low abundance of all histone marks (Fig 2E). Focusing on genes encoding RBFs and ribosomal proteins, revealed that 257 out of 292 expressed genes belong to the third cluster. Examples of genes representative of this cluster include MYC, NAT10, TCOF and FBL (Fig 2F). The colocalization of KDM2B and MYC in the promoters of transcriptionally active genes, suggests that KDM2B, similar to MYC, is a positive regulator of ribosomal biogenesis (36).

Gene Set Enrichment Analyses (GSEA) (37) of the publicly available Cancer Genome Atlas (TCGA) breast cancer (BRCA) dataset, confirmed that the expression of genes involved in ribosome biogenesis is significantly enriched in breast cancers expressing high levels of KDM2B (Normalized enrichment score (NES): 1.81 and q value 0.01) (Fig 3A) and that the enrichment is more robust in basal like BRCA, a breast cancer subtype characterized by enhancement in ribosomal biogenesis (38, 39) (NES 2.54, and q-value <0.001) (Fig 3B). GSEA based on the proteomics data in the CPTAC BRCA dataset, also revealed significant enrichment of proteins involved in ribosome biogenesis in high KDM2B tumors (NES: 2.73 and q value <0.001) (Fig 3C). Additional support to this hypothesis, that KDM2B promotes ribosomal biogenesis in human breast cancer, was provided by the correlations between KDM2B and the KDM2B-regulated genes MYC, UTP14A, TCOF, NAT10 and FBL in the same dataset of basal like breast cancer (Fig 3D) and by the observation that the expression of these genes, similarly to the expression of KDM2B, was highest in the basal-like BRCA subtype (Fig 3E). Importantly, these associations were confirmed in a large independent cohort of triple-negative-breast cancers, published from the Fudan University Shanghai Cancer Center (FUSCC) (40) (Fig 3F and 3G).

**Figure 3.**
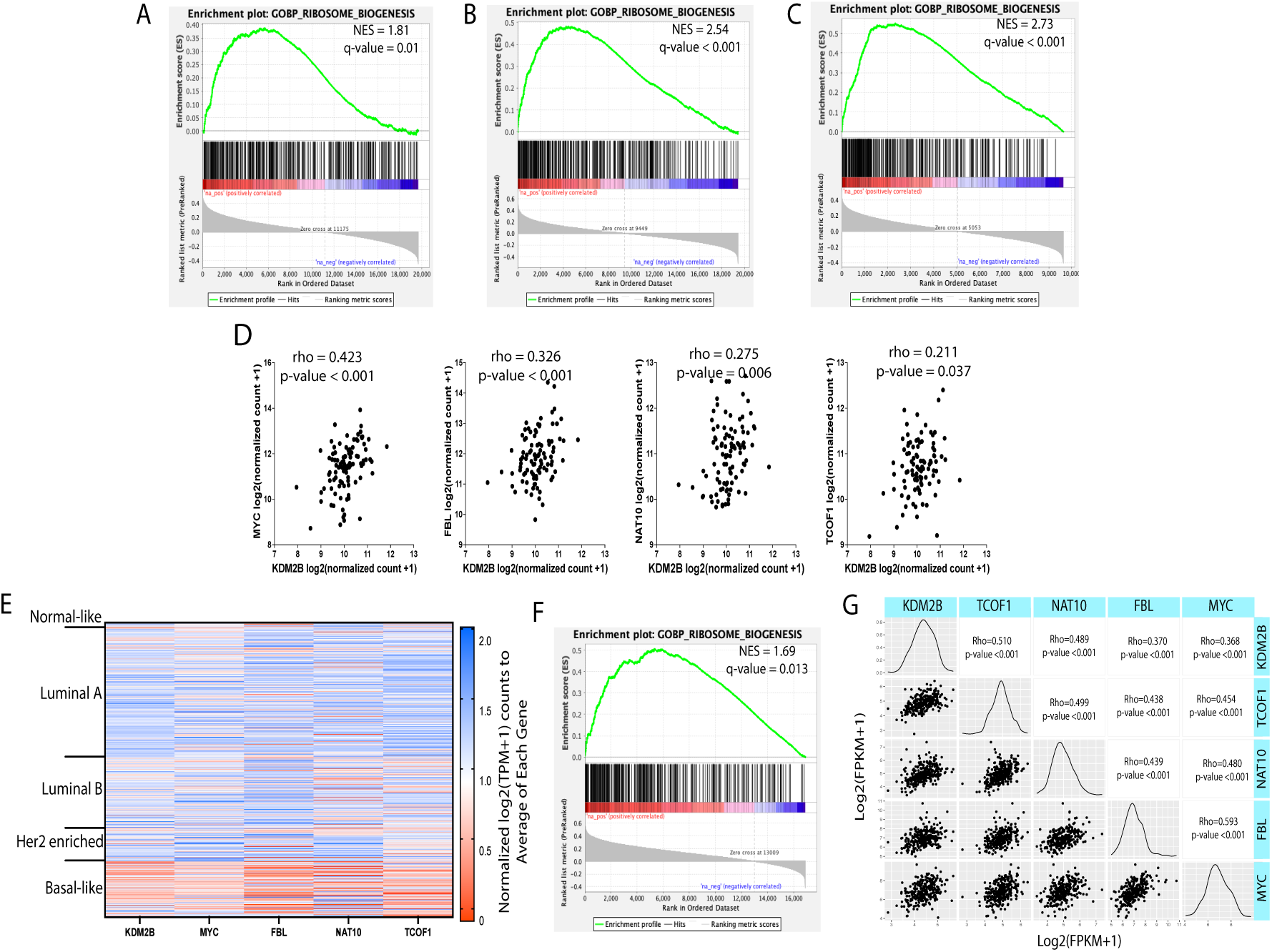
KDM2B expression correlates with the expression of ribosome biogenesis and ribosomal protein-encoding genes in basal-like breast cancer. GSEA of gene sets that were pre-ranked based on the correlation between their mRNA/protein levels and the mRNA/protein levels of KDM2B in the breast cancer TCGA dataset (76) or CPTAC database (78). **A.** Enrichment plot of ribosome biogenesis in the set of all BRCA subtypes in TCGA, **B.** Enrichment plot of ribosome biogenesis in the TCGA basal-like breast cancer subtype only. **C.** Enrichment plot of ribosome biogenesis, generated by GSEA of pre-ranked protein sets from BRCA samples in the CPTAC database (78). **D.** Correlation plots between *KDM2B* and *MYC*, *FBL*, *NAT10*, and *TCOF* in basal-like breast cancer samples in the TCGA database based on PAM50 classifications (37). Correlations and statistics were calculated using the Spearman correlation test on Graphpad Prism. **E.** Heatmap of the relative mRNA levels of *KDM2B*, *MYC*, *FBL*, *NAT10*, and *TCOF1* in the four PAM50 breast cancer subtypes in the TCGA BRCA dataset. The heatmap was made on Graphpad Prism. **F.** GSEA enrichment plot of ribosome biogenesis, generated as described in A, with gene expression data from the set of Triple-Negative BRCAs in the FUSCC patient cohort. **G.** Matrix of correlations plots between *KDM2B*, *MYC*, *FBL*, *NAT10*, and *TCOF1* in the triple-negative BRCA dataset in the FUSCC patient cohort. Correlation plots are displayed on the left, and Spearman rho-values and statistics are displayed on the right. Values displayed are log2(FPKM+1) as calculated by the original authors. Plot made with GGally R package (https://www.rdocumentation.org/packages). ***p-value <0.001.

### The knockdown of KDM2B decreases the steady state levels of ribosomes by lowering the rate of ribosomal biogenesis

The decrease in the abundance of ribosomal proteins likely reflects a decrease in ribosome numbers. To address this hypothesis, we used polysome profiling, which measures the abundance of ribosomal subunits and polysomes. Sucrose density centrifugation of equal amounts of cytoplasmic lysates of control and KDM2B knockdown MDA-MB-231 cells (three independent cultures of each) (Fig 4A), revealed a robust decrease of the free 40S subunits, a smaller decrease of the free 60S subunits, and robust decreases in the 80S ribosomes, and polysomes in the shKDM2B cells (Fig 4B). The same density gradients revealed a larger free ribonucleoprotein peak in the shKDM2B lysates (Fig 4C), suggesting that the knockdown of KDM2B increases the abundance of free, non-ribosome-bound mRNA, a finding consistent with impaired ribosome biogenesis.

**Figure 4.**
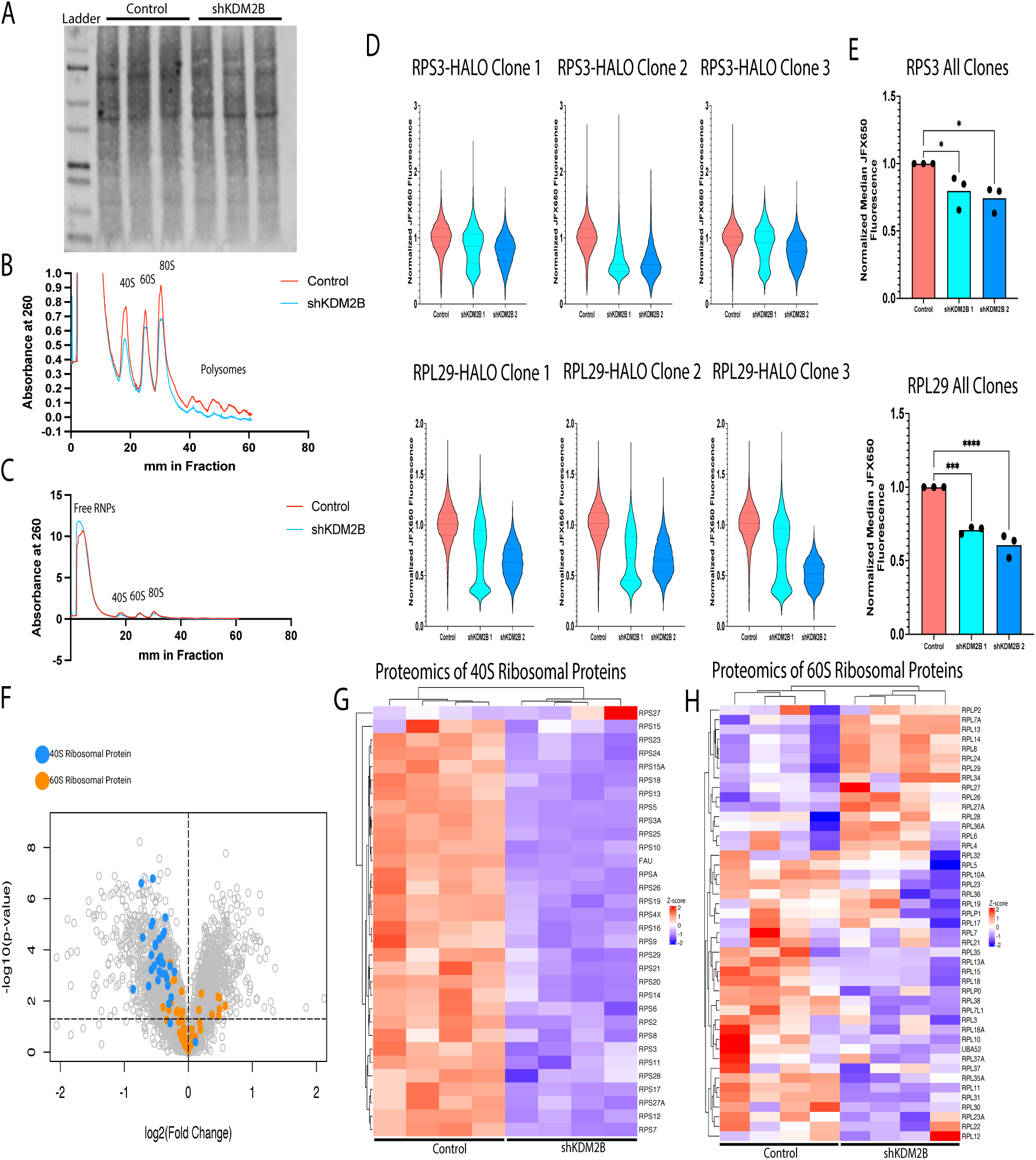
The knockdown of KDM2B reduces ribosome abundance by reducing the rate of ribosome biogenesis. Disproportionate inhibition of the biogenesis of the 40S ribosome. **A.** Protein levels of cytoplasmic lysates derived from control and shKDM2B-transduced MDA-MB-231 cells were quantified using Revert total protein stain. **B.** and **C**. Polysome profiling of equal amounts of the cytoplasmic lysates quantified in A. B shows the absorbance at 260 nm for all the fractions across the gradient, and C focuses on the monosome fractions. Polysome profiling was performed in triplicate, and the display shows the mean signal across all 3 replicates for each condition. **D.** The RPS3 and RPL29 proteins of MDA-MB-231 cells were C-terminally tagged with HALO. Three cell clones with the HALO-tagged RPS3 and three cell clones with the HALO-tagged RPL29 were labeled to saturation with the HALO ligand JFX549. Following the saturation labeling, newly synthesized ribosomes were labeled with the HALO ligand JFX650, and they were analyzed by flow-cytometry 24 hours later. The upper panel shows the distribution of the JFX650 fluorescence intensity relative to the JFX549 fluorescence intensity across individual cells of the RPS3-edited clones, and the lower panel shows the distribution of JFX650 fluorescence intensity relative to the JFX549 fluorescence intensity across individual cells of the RPL29-edited clones. The mean fluorescence intensity (MFI) of the JFX650 signal relative to the JFX549 signal was calculated for RPS3-edited and RPL29-edited cell clones, and each clone had their ratiometric MFI normalized to the control for that clone so that the control was given the value of 1. **E (Upper panel).** The normalized JFX650 MFI ratio of the RPS3-edited control and shKDM2B cell clones. **(Lower panel).** The normalized JFX650 MFI ratio of the RPL29-edited cell clones. Statistics were calculated using a One-way ANOVA, followed by Dunnett’s multiple comparisons test on Graphpad Prism. **F.** Volcano plot showing the log2(fold change) and the – log10(p-value) of the change for all the proteins detected by TMT proteomics in shKDM2B and control cells. Ribosomal proteins are shown as colored dots (blue for the 40S and orange for the 60S proteins). **G** and **H**. Heatmaps showing the relative abundance of 40S (G) and 60S (H) ribosomal proteins, detected by TMT proteomics in control and shKDM2B MDA-MB-231 cells. Unbiased clustering was performed using the ComplexHeatmap R package (https://github.com/jokergoo/ComplexHeatmap). Z-scores were calculated from the geometric mean of the abundance of each protein, and they provided the input for signal intensities, following normalization for each protein. *p-value < 0.05, ***p-value <0.001.

The preceding data suggested a decrease in the abundance of ribosomes in steady state shKDM2B cells. Given that the expression of ribosomal proteins and ribosome biogenesis factors was decreased in shKDM2B cells, we hypothesized that the decrease in the number of ribosomes was due to a decrease in the rate of ribosome biogenesis. To address this hypothesis, we took advantage of the fact that free ribosomal proteins have short half-lives, but they are stabilized by association with the ribosomes, whose half-life is exceedingly long (41–43). Therefore, the accumulation of such proteins over time is proportional to the rate of ribosome assembly, and the time-dependent change of their abundance provides a surrogate measure of the rate of ribosome biogenesis. To monitor newly synthesized ribosomal proteins, we first tagged the endogenous RPL29 and RPS3 proteins in MDA-MB-231 cells with a C-terminal Halo tag, using CRISPR-Cas9-based gene editing (44) (Fig S2A and S2B). RPL29– and RPS3-edited, control and shKDM2B cells were then treated to saturation for one hour with a tenfold molar excess, based on protein molar calculations in OpenCell (45), of the Halo-tag ligand JFX549 (excitation 549 nm/emission 570 nm) (46, 47) that covalently and irreversibly (48) binds the Halo-tag. Following washing to remove the excess ligand, newly synthesized ribosomes were pulse-labeled with JFX650, a second Halo-tag ligand (excitation 650 nm/emission 667 nm). Tracking the JFX650 fluorescence 24 hours later by flow cytometry, showed that the normalized median JFX650 fluorescence was consistently decreased in all shKDM2B cells (Fig 4D and 4E).

Polysome profiling (Fig. 4B) suggested that there may be an imbalance between the 40S and 60S ribosomal subunits in KDM2B knockdown cells. Strong support to this hypothesis was provided by the shKDM2B-induced downregulation of ribosomal proteins, which was significantly more consistent and robust for the 40S than the 60S proteins (Fig 4F, 4G and 4H).

### The knockdown of KDM2B gives rise to defects in pre-ribosomal RNA processing

The assembly and maturation of ribosomal subunits occurs in the nucleolus and depends on the recruitment of small nucleolar RNAs (snoRNAs), and large sets of ribosome biogenesis factors. Ribonucleoproteins composed of snoRNAs, and RBFs direct the cleavage of the 47S pre-ribosomal RNA, which is controlled by precisely orchestrated RNA and protein movements (14, 15, 18, 27, 49, 50). Given that the abundance of many of these RBFs is significantly downregulated by the knockdown of KDM2B (Fig 1C and 1D), we hypothesized that KDM2B depletion would impair pre-ribosomal RNA processing, with perhaps stronger effects on the processing events leading to the 18S RNA, a component of the more strongly impacted 40S ribosomal subunits. Additionally, given that the maturation of ribosomal subunits in the small subunit (SSU) and large subunit (LSU) processomes is precisely orchestrated, identification of the cleavage events primarily affected by KDM2B depletion, should inform the stage of maturation critically targeted by the knockdown of KDM2B.

The 47S RNA contains in addition to the sequences of the mature ribosomal RNAs, two External Transcribed Spacers (5′ and 3′ ETS) and two Internal Transcribed Spacers (ITS1 and ITS2). The spacers are removed during processing, and they are not part of the highly abundant mature ribosomal RNAs (Fig S3). Therefore, spacer sequences can be used to probe Northern blots monitoring ribosomal RNA processing. The first pre-ribosomal RNA processing step consists of two cleavage events, which remove the 3′ ETS (02 cleavage) and a short fragment of the 5′ ETS (A′ cleavage) from the 47S rRNA, giving rise to the 45S rRNA. The latter is cleaved at an ITS1 site upstream of 5.8S (cleavage 2), giving rise to the 30S and 32S rRNAs, which are precursors of the mature 18S and 28S rRNAs respectively. The 30S rRNA undergoes two cleavage events (A0 and A1) which remove the 5′ ETS, giving rise to the 26S and 21S precursors of the mature 18S rRNA. Alternatively, these two cleavage events may occur in the 45S rRNA, prior to cleavage 2, giving rise to 43S and 41S rRNAs. These, and all the remaining rRNA processing steps are shown in Figure S3.

To determine whether KDM2B depletion results in pre-ribosomal RNA processing defects, we first probed a Northern blot of total RNA from three control and three shKDM2B cultures of MDA-MB-231 cells with the ITS1 probe (Fig S3). This probe can theoretically detect the 47S/45S, 43S, 41S and 30S rRNAs, as well as the 30S-derived precursors of the mature 18S rRNA. The results showed that whereas the 30S rRNA was more abundant in shKDM2B cells, the abundance of the 26S and 21S rRNA species was significantly lower (Fig 5A and 5B). This observation suggests that the efficiency of the A0 cleavage in the 5′ ETS of the 30S rRNA, which generates the 26S rRNA, is impaired in the KDM2B knockdown cells. Interestingly, whereas the ratio of 26S to 30S rRNA was significantly lower in shKDM2B cells (Fig 5C, left panel), the ratio of the 26S to 21S rRNA, was unchanged (Fig 5C, right panel), suggesting that a KDM2B-dependent pathway regulates the A0 cleavage, and that the A1 cleavage, proceeds normally once the A0 cleavage has occurred. Two significantly less abundant species migrating slower than the 30S rRNA may be the 43S and 41S. Their low abundance relative to the 30S rRNA suggests that the first 45S rRNA site to be cleaved in MDA-MB-231 cells is site 2, the same site shown earlier to be the first one cleaved in Hela cells (51). A second experiment, probing the same RNAs with the 5′ ETS probe confirmed the slightly higher abundance of the 30S rRNA and its defective processing to the 26S rRNA (Fig 5D and 5E). In this experiment we did not detect the 21S rRNA because the 5′ ETS probe cannot detect this RNA (Fig S3).

**Figure 5.**
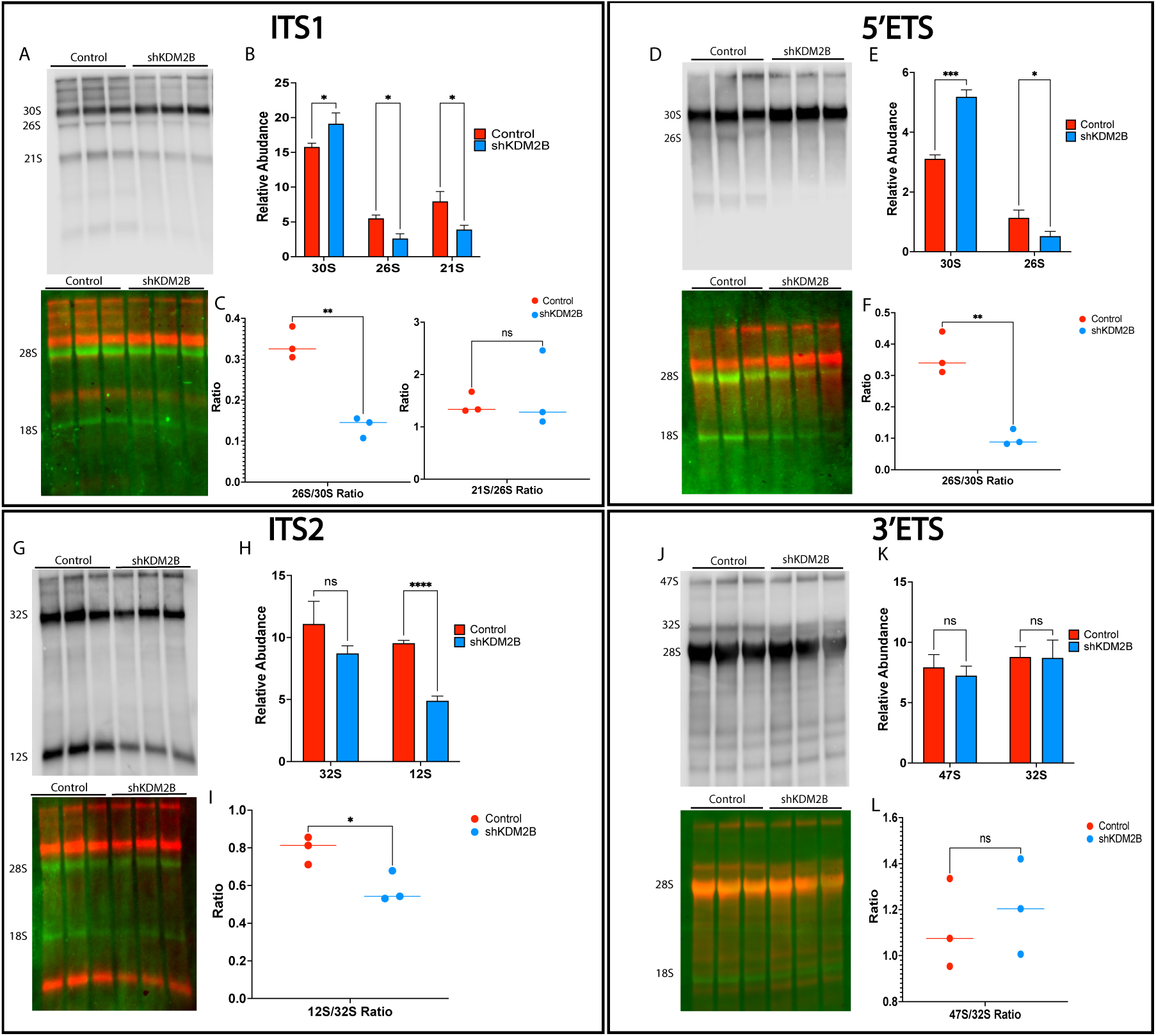
The knockdown of KDM2B impacts pre-ribosomal RNA processing. Northern blots of pre-ribosomal RNA isolated from three independent cultures of control or shKDM2B-transduced MDA-MB-231 cells were probed with digoxigenin (DIG)-labeled single strand RNA probes of ITS1 (A, B and C), 5′ ETS (D E, and F), ITS2 (G, H, and I) and 3′ ETS (J, K, and L) (see supplementary figure S3 for the position of the probes within the map of the 47S pre-ribosomal RNA). Hybridization was detected with an alkaline-phosphatase tagged anti-DIG antibody and chemiluminescence. The upper panels in A, D, G, and J show the chemiluminescence signal, and the lower color panels show the 18 and 28S RNAs, which are labeled with Ethidium bromide (600 nm pseudo-colored in Green) and the rRNA intermediates (chemiluminescence signal pseudo-colored in Red). B and C, E and F, H and I and K and L show the quantification of the rRNA intermediates for panels A, D, G, and J, respectively. Statistics were calculated using the multiple unpaired t-test and correcting for multiple hypothesis testing with the Holm-Šídák method on Graphpad Prism. Error bars represent the standard deviation from the mean. *p-value < 0.05, **p-value <0.01, ***p-value <0.001, ns, not significant.

The preceding experiments confirmed that KDM2B promotes the processing of 30S rRNA, a precursor of the 18S rRNA. To determine whether the pre-ribosomal RNA processing leading to the mature 28S and 5.8S rRNAs is also under the control of KDM2B, we probed the same RNAs with the ITS2 probe (Fig S3), which should detect the 32S rRNA and its downstream product 12S, the precursor of the 5.8S rRNA. The results showed that both the 32S and 12S rRNAs were less abundant in shKDM2B cells (Fig 5G and 5H). However, the downregulation of the 12S rRNA was slightly more robust (Fig 5H), resulting in the modest lowering of the ratio 12S/32S (Fig 5I), and suggesting that cleavage 4, which generates 12S from the 32S rRNA, was also impaired. The low abundance of the 30S rRNA could be the result of a partial defect in cleavage 2, which is catalyzed by UTP24 and gives rise to the 30S and 32S pre-ribosomal RNAs. Alternatively, the loss of KDM2B may destabilize the 32S rRNA. We propose that the instability of 32S is a more likely explanation, as it was also suggested by the Northern blot (Fig. 5J), which was probed with the 3′ ETS probe, and which showed that the ratio 47S/32S was not affected by the KDM2B knockdown (Fig 5L), and that the 32S rRNA was detected as a smear and not as a distinct band. Although the loss of KDM2B affects the pre-ribosomal RNA processing events that give rise to both the 18S and the 28S rRNA, its effects on the processing events leading to the 18S rRNA were significantly more robust, in agreement with the polysome profiling and quantitative proteomics data (Fig 4). The proposed destabilization of the 32S rRNA may be a compensatory cellular mechanism that aims to minimize the imbalance. This agrees with recent studies showing that rRNA not yet incorporated into the assembled pre-ribosomal units, may undergo PAPD5-dependent polyadenylation, followed by 3′-5′ degradation by the exosome (52).

The transcriptomic and quantitative proteomic data in figure S4, confirm that the expression of ribosome biogenesis factors associated with the SSU processome is significantly impacted by the depletion of KDM2B, but they do not address how the observed changes impact the maturation of the SSU processome. This will be addressed in future studies, by integrating functional KDM2B knockdown data, with data on the structural dynamics of processome assembly and maturation (14, 15, 18, 27, 49, 50).

### The knockdown of KDM2B decreases global mRNA translation

The observed defect in the biogenesis of the 40S ribosomal subunit, suggested that the knockdown of KDM2B may decrease the global rate of mRNA translation. To address this hypothesis, we pulsed methionine-starved control and KDM2B knockdown MDA-MB-231 cells with L-Homopropargylglycine (L-HPG), an amino-acid analog of methionine that has been modified with an alkyne group. Nascent proteins labeled with the modified amino acid over time were detected with biotin-azide via “click” chemistry. This showed that the knockdown of KDM2B results in a 66% decrease of the global rate of translation. While L-HPG incorporation was detected in control cells at 7.5 minutes, it was first detected in shKDM2B cells at 30 minutes. Moreover, L-HPG incorporation in KDM2B knockdown cells lagged behind compared to control cells for the entire 3-hour period of monitoring (Fig 6A). Quantification of the data in Figure 6A revealed a significant drop in the rate constant of L-HPG incorporation over time in KDM2B knockdown, relative to the control cells (0.2937 Arbitrary Units (AU) vs 0.1295 AU) (Fig 6B). mRNA translation was reduced at all time points by approximately 60-70% (Fig 6B and 6C).

**Figure 6.**
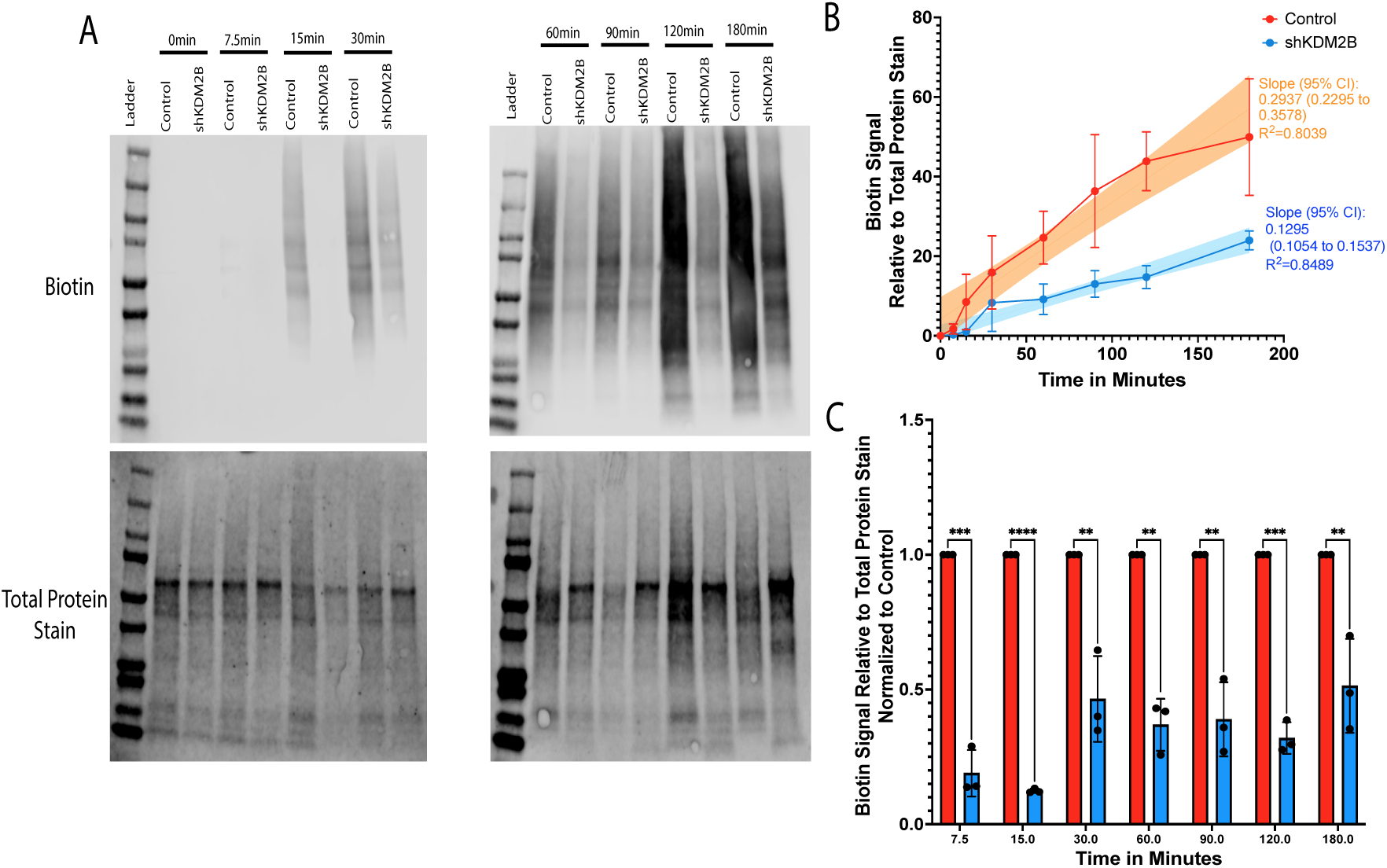
The knockdown of KDM2B reduces the global rate of mRNA translation. **A.** Methionine starved control and shKDM2B-transduced MDA-MB-231 cells were pulsed with the methionine analog L-HPG. Cell lysates harvested at the indicated time points were electrophoresed and transferred to PVDF membranes. (Upper panels) Newly synthesized proteins were detected and quantified with biotin-azide via click chemistry. (Lower panels) Protein loading per lane was visualized and quantified with Revert Protein Stain. The experiment was done on three biological replicates, and the results were similar in all three. Panels show one of the replicates as a representative example. **B.** Biotin signal normalized to the total protein signal for all replicates in each time point and condition. Simple linear regression with line of best fit calculation was performed to determine the relative rates of protein production in arbitrary units. The line of best fit was plotted with the 95% confidence interval displayed as an orange or blue backdrop in the two data series. The plot was generated on Graphpad Prism. **C.** Quantification of the biotin signal in the three shKDM2B replicates relative to the three control replicates at each time point. Statistics were calculated using the multiple unpaired t-test and correcting for multiple hypothesis testing with the Holm-Šídák method on Graphpad Prism. Error bars represent the standard deviation from the mean. **p-value <0.01, ***p-value <0.001, ****p-value <0.0001.

### The knockdown of KDM2B relieves ISR in cancer cells expressing high levels of KDM2B

The decrease in the rate of the global mRNA translation in shKDM2B cells is likely to be caused by the low abundance of ribosomes. Another factor that may contribute to the translational repression is the activation of stress response mechanisms. One of these mechanisms is the integrated stress response (ISR), which can be activated by amino acid deprivation occurring in KDM2B knockdown cells because of defects in the biosynthesis of multiple non-essential amino acids(12). The main hallmark of ISR activation is the phosphorylation of eIF2α at Ser51, which is catalyzed by several eIF2α kinases. Phosphorylation of eIF2α increases its affinity to eIF2B and interferes with the eIF2B guanine nucleotide exchange activity (53). This inhibits the formation of the eIF2-GTP-tRNA (Met) ternary complex and translational initiation (54). However, contrary to expectation, the phosphorylation of eIF2α was reduced in KDM2B knockdown cells (Fig S5A), along with the expression of the eIF2α kinases PERK and PKR (Fig S5B). Importantly, the expression of MYC, a known activator of the eIF2α kinase GCN2 (55), was also reduced in the same cells (Fig 1). We conclude that ISR is not a contributing factor to the shKDM2B-induced translational repression.

### The knockdown of KDM2B dysregulates transcript specific translation

To explore how the knockdown of KDM2B impacts the dynamics of mRNA translation we performed ribosome profiling (RIBO-Seq) in control and KDM2B knockdown MDA-MB-231 cells. The quality of the RIBO-Seq data, was addressed by examining the trinucleotide periodicity of ribosome protected fragments along the entire length of all transcripts. A prerequisite to addressing the periodicity is the accurate mapping of the ribosomal peptidyl site (P-site) and the determination of the distance of this site from the 5′ and 3′ ends of ribosome protected fragments of different lengths (P-site offset, PO). This was done using riboWaltz, a bioinformatics tool implemented in R, and available as an R package. RiboWaltz analyses revealed that most of the signals map in mRNA coding regions (Fig S6A) and display clear trinucleotide periodicity (Fig S6B and S6C), confirming the quality of the data. Mapping the distribution of ribosome protected fragments across the length of mRNAs genome wide, revealed reduced signals in the start and stop codons in shKDM2B cells (Fig S6B and S6C), which is consistent with reduced global translation, due to the lower abundance of ribosomes in these cells (Fig 4B).

Pairing the RIBO-seq (56) data, with data derived from parallel RNA-Seq studies, allowed us to measure the difference in mRNA translational efficiency (ΔTE) between control and shKDM2B cells. Comparisons of the ΔTE values in the two cell types revealed that for some transcripts the ΔTE was not affected by the knockdown of KDM2B, while for other transcripts it was either decreased or increased (Fig S6E). This analysis confirmed that the shKDM2B-induced global translational repression is distributed unevenly among mRNA transcripts. Differences in transcript specific translation could be the result of codon bias. This hypothesis was supported by our observation that the knockdown of KDM2B resulted in widespread changes in amino acid metabolism (12), as well as in reduced tRNA amino acyltransferase expression (Fig S7A and S7B). Plotting the P-site signal of all codons in control, versus shKDM2B cells however, failed to reveal statistically significant KDM2B-dependent differences (Fig S7C and S7D), suggesting that dysregulation of transcript-specific translation was not due to codon bias.

Another potential cause of transcript-specific dysregulation is ribosome heterogeneity (57). The shKDM2B-induced changes in the expression of ribosomal proteins varied in strength (Fig 4F), suggesting that KDM2B may not only impact ribosome abundance, but also ribosome composition. To address this question, we employed a recently published strategy, which uses the abundance of ribosome protected rRNA fragments to predict the heterogeneity of ribosome-incorporated ribosomal proteins in control and shKDM2B cells (58). This analysis revealed that the knockdown of KDM2B does not have a robust ribosome heterogeneity signature (Fig S8). Only two ribosomal proteins, RPS4X (eS4) and RPS8 (eS8), were significantly enriched. However, their enrichment was weak, raising questions whether it reflects true ribosomal heterogeneity.

The efficiency of mRNA translation has been shown to depend on CDS and/or mRNA length (56, 59, 60). We therefore questioned whether transcript and CDS length influence the transcript sensitivity to the shKDM2B-induced translational repression. Analysis of the RIBO-Seq data in control and shKDM2B-tansduced cells revealed that whereas transcript length does not affect translational efficiency (Fig 7A), CDS length has an impact, with long CDS transcripts being translated more efficiently than short CDS transcripts in shKDM2B cells (Fig 7B). Translation efficiency is also known to depend on structural motifs and upstream ORFs (uORFs) that are located primarily in the 5′ UTR (22, 61, 62). The analysis of the RIBO-Seq data showed that whereas the translational efficiency of transcripts with short unstructured, AU-rich 5′ UTRs tends to be reduced in shKDM2B cells (Fig 7C, 7E, and 7F), the translational efficiency of transcripts, with long structured, GC rich 5′ UTRs tends to remain unchanged, or to increase (Fig 7D, 7G and 7H). In agreement with these data, *de novo* motif analysis (STREME), comparing the 5′ UTR of transcripts with decreased or increased translational efficiency to the 5′ UTR of transcripts whose translational efficiency was unchanged revealed that transcripts with decreased translational efficiency were enriched in uracil-rich motifs whereas those with increased translationally efficiency were enriched in cytosine-rich motifs (Fig S9A and S9B). Similarly, Analysis of Motif Enrichment (AME) revealed that the top five most enriched motifs in the 5’ UTR of mRNAs whose translational efficiency was increased in shKDM2B cells, were with few exceptions, GC rich while the top five most enriched motifs of mRNAs whose translational efficiency was decreased, were AU-rich (Fig S9C and S9D). Another factor that could potentially influence the ΔTE of different mRNAs in an environment of decreasing ribosome numbers, is mRNA abundance. Low abundance may reduce the ability of mRNAs to compete when translational initiation complexes are reduced, as is the case in KDM2B knockdown cells. However, this is not the case. In fact, the impact of the depletion of KDM2B on the translational efficiency of high abundance mRNAs is significantly more robust, than the impact of KDM2B depletion on low abundance mRNAs (Fig 7I).

**Figure 7.**
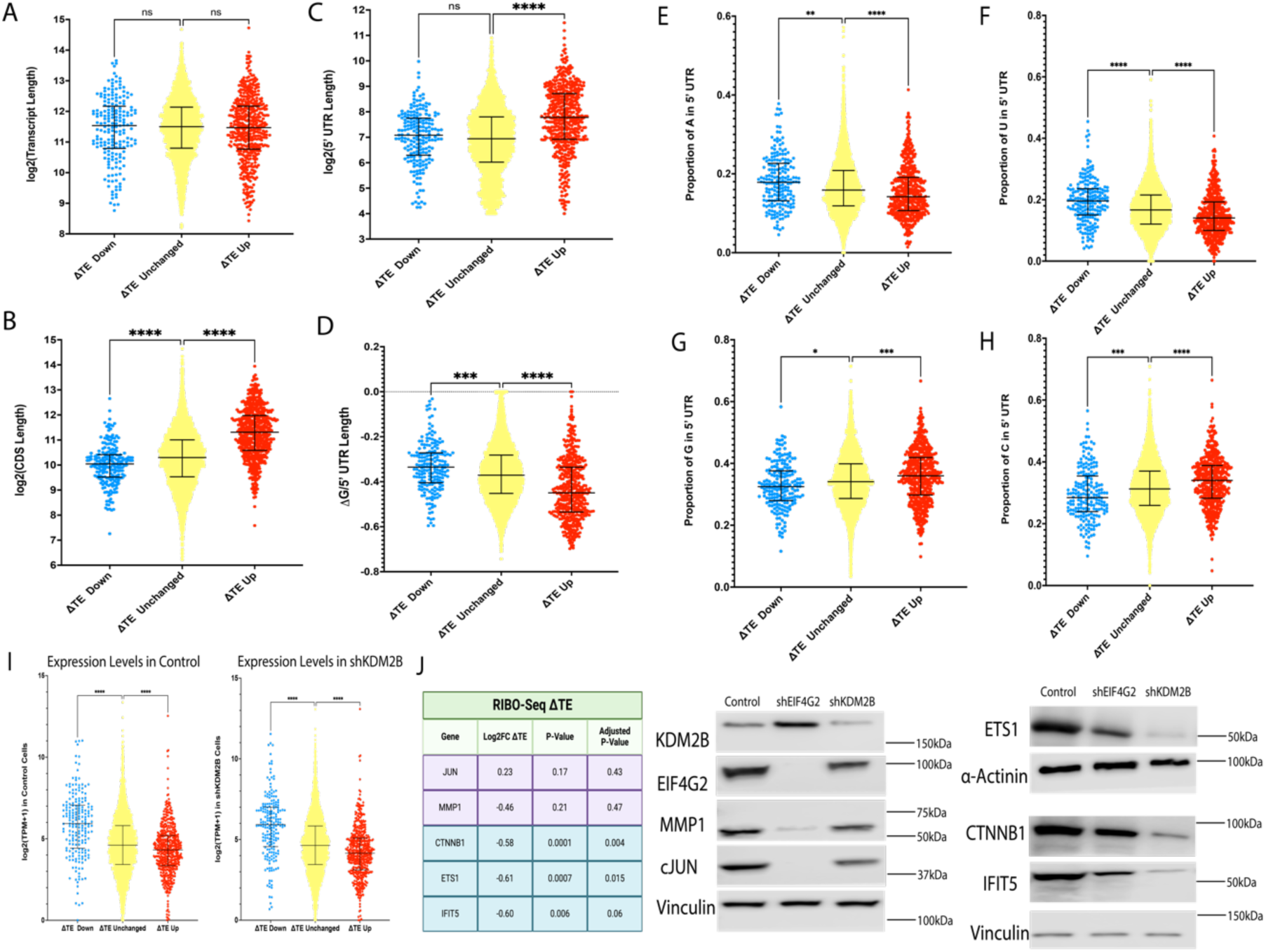
The knockdown of KDM2B impacts transcript-specific translation. Transcripts, which in shKDM2B-transduced cells exhibit significantly decreased translational efficiency (log2(FC)<-0.5), increased translational efficiency (log2(FC)>0.5), or unchanged translational efficiency, were plotted for **A.** Total Transcript length, **B.** CDS length, **C.** 5′ UTR length, **D.** 5′ UTR free energy (ΔG), **E**. 5′ UTR adenine composition, **F**. 5′ UTR uracil composition, **G**. 5′ UTR guanosine composition, **H**. 5′ UTR cytosine composition, and **I.** Total Transcripts levels in transcripts per million (TPM) in control and shKDM2B-transduced cells. Statistics were calculated with the Kruskal-Wallis test followed by a Dunn’s multiple comparisons test. **J.** Table of translational efficiency of select transcripts, followed by western blotting comparing the abundance of the proteins encoded by these transcripts in shKDM2B and shEIF4G2 cells. *p-value < 0.05, **p-value <0.01, ***p-value <0.001.

The translation of some mRNAs with long 5′ UTRs that are predicted to have robust secondary structures, has been shown to be regulated by alternate mechanisms of translation initiation, raising the question of the role of KDM2B in the regulation of these mechanisms. Alternate mechanisms of translation initiation depend on complexes other than the canonical eIF4F complex, which is composed of the scaffold protein eIF4G, the cap binding protein eIF4E and the RNA helicase eIF4A. The known cap binding proteins (CBPs) of these alternative complexes include eIF3d(63), NCBP2/CBP20 (64, 65), and NCBP3 (65). RNA binding scaffold proteins recruited by these CBPs include the eIF4GI homolog eIF4G2 (also known as DAP5, NAT1, or p97), which lacks the eIF4E binding domain and utilizes the cap binding protein eIF3d instead (66). Confirmed targets of the eIF4G2/eIF3d complex include c-JUN and MMP1(66). Importantly, the ΔTE of these mRNAs in shKDM2B relative to the control MDA-MB-231 cells was not statistically significant (Fig 7I left panel), suggesting that their translation may indeed be insensitive to the loss of KDM2B. Addressing the expression of the proteins encoded by these mRNAs in control, shKDM2B and sheIF4G2-transduced cells, confirmed that whereas shKDM2B had only a minimal effect (Fig 7J middle panel), shIF4G2 inhibited their expression profoundly. The expression of the proteins encoded by the mRNAs of CTNNB1, ETS1 and IFIT5, whose translational efficiency was strongly inhibited in shKDM2B cells on the other hand, was significantly downregulated by shKDM2B and only minimally affected by sheIF4G2 (Fig 7J right panel).

These data combined, confirmed that the translational repression induced by the knockdown of KDM2B is unevenly distributed among mRNA transcripts and provided evidence that some of the mRNAs whose translation is insensitive to the loss of KDM2B are regulated by alternative mechanisms of translational initiation. Given that the knockdown of KDM2B inhibits global mRNA translation, mRNAs with long CDS and/or complex GC-rich 5′ UTRs and ΔTE upregulation likely represent mRNAs whose translational repression may be less pronounced than that of mRNAs with short CDS and less complex 5′ UTRs.

### KDM2B coordinates mRNA translation with cell cycle progression

Evidence presented in this report shows that the knockdown of KDM2B, a proliferation-promoting oncoprotein, inhibits mRNA translation, as expected. Moreover, Metascape enrichment analysis (67) based on all transcripts whose translational efficiency was scored in our paired RIBO-Seq/RNA-Seq study, showed that the most enriched among the functions regulated by mRNAs with reduced translational efficiency in KDM2B knockdown cells is the cell cycle (Fig S10A). Given the interdependence of mRNA translation and the cell cycle, KDM2B may regulate translation by targeting the cell cycle, or vice versa. Alternatively, it may regulate both independently. In either case, it functions as a master integrator of two functions, whose activity needs to be coordinately regulated.

Earlier observations had shown that the efficiency of mRNA translation fluctuates during the cell cycle, with robust translational efficiency during the G1 phase, and translational repression during mitosis (68, 69). Given that global translational efficiency is repressed in shKDM2B cells (Fig 6), despite the fact that they accumulate in G1 (Fig S10B), a phase characterized by robust mRNA translation, we conclude that the shKDM2B-induced translational repression is most likely the primary event. This is also supported by the data in the volcano plot of the results of our paired RIBO-Seq/RNA-Seq experiment, which show that shKDM2B decreases selectively the translational efficiency (ΔTE down) of most mRNAs encoding proteins that promote the transition from G1 to S (color-coded in purple in Fig S10C). These data collectively suggest that KDM2B coordinates the activities of mRNA translation and the cell cycle, either by targeting translation upstream of the cell cycle, or by targeting both simultaneously.

## DISCUSSION

Earlier studies had suggested that KDM2B encodes a nucleolar protein which inhibits ribosomal biogenesis by repressing the transcription of pre-ribosomal RNA (26). In this report we revisited this unexpected finding in control and KDM2B knockdown MDA-MB-231 cells, using an unbiased multiomics strategy. The results unequivocally showed that KDM2B promotes, rather than inhibiting ribosomal biogenesis and mRNA translation. First, it promotes the expression of many ribosome biogenesis factors, and ribosomal proteins with its effects being more robust on the expression of factors involved in the assembly of the small subunit (SSU) processome, and on the expression of proteins associated with the 40S ribosomal subunits. Consistent with this, the knockdown of KDM2B was also shown to inhibit the processing of pre-ribosomal RNA, with robust inhibition of the A0 cleavage in the 5′ ETS, and weak inhibition of cleavage 4, in ITS2. The A0 cleavage occurs during the transition of the SSU processome from the pre-A1 to the post-A1 stage (27). The impaired ribosome biogenesis results in global inhibition of mRNA translation. However, coupled RIBO-Seq and RNA-Seq studies in the same cells, also showed that ribosomal changes induced by the knockdown of KDM2B modulate unequally the efficiency of translation of different mRNAs, inhibiting more robustly the translation of high abundance mRNAs with a short CDS and with short unstructured 5′ UTRs. The global and transcript-specific effects on mRNA translation appear to be caused by the low abundance of ribosomes in KDM2B knockdown cells, as the potential contribution of ISR, ribosome heterogeneity and codon bias was excluded.

The most detailed studies in ribosomal biogenesis have been carried out in the yeast *Saccharomyces cerevisiae.* However, recent cryo-EM studies of the human SSU processome provided excellent structural information on the events associated with the early steps of ribosome biogenesis in human cells as well. These studies identified three structurally distinct states: Pre-A1, pre-A1* and post A1(27, 70). The 5′ ETS is present in the pre-A1 and pre-A1* states. In the post-A1 state, only a small segment of the 5′ ETS remains between the interacting UTPA and UTPB complexes. The first step in the process of maturation is the release of Neuroguidin (NGDN), which results in major conformational changes of multiple ribosome assembly factors and ribosomal proteins, including HEATR1 (UTP10), UTP20, RPS4x (eS4), RPS9 (uS4) and U3IP2 (RRP9). These changes, which mark the transition from pre-A1 to pre-A1*, create a vacant space, which is occupied by the disordered helix 21 of the 18S rRNA. During the transition from the pre-A1* to the post-A1 stage, five factors (NAT10, NOL10, AATF, KRR1 and C1orf131) are released and five additional factors (DHX37, DIM1, ExosC10, AROS and RPS19) are recruited. Our data do not address how the changes induced by the knockdown of KDM2B impact the exquisite choreography of these events. However, they do show that KDM2B depletion impairs the efficiency of the A0 cleavage in the 5′ ETS, which suggests a defect in the maturation of the SSU processome around stage A*. They also show that KDM2B depletion impairs weakly the efficiency of cleavage 4, in ITS2, suggesting an additional stage-specific defect in the maturation of the LSU processome. Identification of the ribosomal biogenesis stages impacted by regulatory factors like KDM2B, is a prerequisite for future studies that will address the dynamic regulation of the process in all its complexity.

The inhibition of ribosome biogenesis induced by the knockdown of KDM2B decreases ribosome abundance and inhibits global mRNA translation. However, coupled RIBO-seq and RNA-Seq data also showed that ribosomal changes induced by the knockdown of KDM2B alter unequally the efficiency of translation of different mRNAs, inhibiting more robustly the translation of mRNAs with short CDS, and/or short unstructured 5′ UTRs. Data in this report linked the inhibition of ribosome biogenesis in shKDM2B cells, to the selective inhibition of translation of short mRNAs, a finding reminiscent of earlier observations showing that the translational efficiency of short transcripts is more sensitive to the depletion of ribosome biogenesis factors than the translational efficiency of long transcripts (56, 59, 71–74). Interestingly, transcripts with short CDS and/or short unstructured 5′ UTRs, tend to exhibit higher ribosome occupancy and basal translational efficiency (56, 59). This suggests that these transcripts recruit ribosomes more efficiently and explains why they may be more sensitive to changes in ribosome abundance as they may be dependent on higher ribosomal abundance to achieve high translational efficiency. The differences in translation between transcripts could potentially be explained by the closed loop model of translation (75), which suggests that mRNAs undergoing translation circularize, bridging their 5′ and 3′ ends, which facilitates translational re-initiation. Mathematical modeling of this concept, using yeast or human data, predicted that transcripts with a short CDS re-initiate more efficiently and exhibit higher ribosome density, perhaps by forming closed loops more efficiently than transcripts with a long CDS (60), and this prediction has been experimentally confirmed (72–74).

The preceding data suggest that shKDM2B may inhibit unequally the translation of short and long mRNA transcripts, simply by affecting the abundance of ribosomes. However, additional mechanisms may also contribute to this outcome. One of these mechanisms depends on RACK1 (72–74), a protein which associates with the 40S ribosomal subunit, and which is significantly downregulated in KDM2B knockdown cells (this report). RACK1 mutants inhibit selectively the translational initiation of closed loop forming short mRNA transcripts perhaps by inhibiting their circularization(72–74). Another potential mechanism depends on the translational initiation complex that is preferentially used for the translation of different mRNAs. The canonical translational initiation complex eIF4F consists of the cap-binding protein eIF4E, the scaffold protein eIF4G1 and the RNA helicase eIF4A. However, mRNAs with a long CDS and highly structured 5’UTRs, which tend to be less sensitive to changes in the expression of KDM2B, may employ alternate translation initiation complexes that do not contain eIF4G1. Importantly, whereas eIF4G1 interacts with the polyA binding protein (PABP) and facilitates mRNA circularization, the scaffold proteins employed by the alternate translational initiation complexes do not. As a result, such transcripts circularize less efficiently, preventing the occurrence of sequential uninterrupted cycles of translation. Data presented in this report confirmed the relative insensitivity of some mRNAs which employ preferentially an eIF4G2-containing alternate translational initiation complex, to the KDM2B knockdown.

The differential interpretation of changes in ribosome number by different mRNAs in KDM2B knockdown cells may have significant implications on our understanding of the role of KDM2B in cells responding to different cell autonomous and environmental signals. Translation of different mRNAs is not exclusively under the control of one mechanism. Instead, it may depend on combinations of alternate, along with the canonical mechanism of translational initiation in different settings. What determines the preferred mechanism of initiation is unknown. Also, it is unknown whether the different choices depend on cell type and tissue specific regulatory mechanisms, which can be modulated by different cell intrinsic or environmental signals. Perhaps, conditions like stress, drug resistance viral infection and transformation may reprogram the cells via KDM2B and other factors, altering ribosome abundance, ribosomal composition, and the corresponding preferred complex of translational initiation. The ultimate outcome of the type of complex reprogramming we propose may be the emergence of signal and transcript-specific changes in the efficiency of translation. Our data suggest that KDM2B may be an important regulator of the proposed reprogramming.

## MATERIALS AND METHODS

Detailed materials and methods can be found in the Supplementary Materials and Methods. A brief summary of the materials and methods is presented.

### Cells and Culture Conditions

MDA-MB-231 and HEK293T-LentiX cells were cultured in High Glucose Dulbecco’s modified Eagle’s minimal essential medium, supplemented with 10% fetal bovine serum, penicillin, streptomycin, sodium pyruvate, non-essential amino acids, and Plasmocin™. Cells were subcultured every 2-3 days, with identity validated via Short Tandem Repeat (STR) profiling with annual re-validation. Regular mycoplasma testing was conducted using a PCR-based assay. Plasmid transfection was performed using Lipofectamine 3000™ or electroporation as detailed in the Supplemental Materials and Methods. Cell cycle distribution was analyzed using propidium iodide staining and flow cytometry, with data analyzed on FlowJo. Lentiviral packaging was performed using Lipofectamine-based transfection with 2nd generation packaging plasmids into Lenti-X cells, followed by virus collection, filtration, and transduction.

### Molecular Profiling

Transcriptomics using RNA-seq, proteomics using Tandem Mass Tag mass spectrometry, and translatomics using paired RIBO-seq/RNA-seq were employed to compare control and shKDM2B transduced MDA-MB-231 cells. Chromatin binding of KDM2B, MYC, H3K4me3, H3K27me3, H3K27ac, and H2AK119ub was assessed using ChIP-seq. Chromatin accessibility was assessed using ATAC-seq comparing control and shKDM2B transduced MDA-MB-231 cells.

For immunoblotting, cells were lysed in RIPA buffer with protease and phosphatase inhibitors, followed by protein concentration determination using the Pierce™ BCA Protein Assay Kit. Proteins were separated by electrophoresis, transferred to PVDF membranes, and probed with specific antibodies. Membranes were imaged using a LI-COR Odyssey® Fc Imaging System.

### Biochemical Assays

For tracking of translation rates, cells were starved of methionine, cystine, glutamine and then grown in L-homopropargylglycine (L-HPG). Cells were collected and lysed, and newly synthesized proteins containing L-HPG were labeled with biotin using click-chemistry. Labeled proteins were analyzed by electrophoresis and immunoblotting.

To assess ribosome abundance, Polysome profiling was performed via sucrose gradient ultracentrifugation of cytoplasmic lysates. Gradient fractions were analyzed by absorbance at 260 nm. Additionally, CRISPR editing of the RPS3 and RPL29 loci with Halo allowed the tracking of ribosome production (44).

The processing of pre-ribosomal RNA in control and shKDM2B-transduced cells, was monitored by Northern blotting. To detect different pre-ribosomal RNA processing intermediates, RNA was extracted, electrophoresed, transferred to membranes, and probed with DIG-labeled antisense probes. Membranes were imaged for chemiluminescence signals emitted by the hybridized probes.

### Patient Databases

Gene expression data from TCGA (76), GTEx (77), and FUSCC (40) datasets were obtained and analyzed for correlations between ribosome biogenesis factors and KDM2B. Proteomic data from CPTAC (78) were also analyzed for correlations between ribosome biogenesis factors and KDM2B.

## Author Contributions

VA and PNT conceived the project, designed experiments, and wrote the manuscript. VA performed most of the experiments and analyzed data. EC performed experiments and analyzed data. ALF carried out bioinformatic analyses with support from IC and GN. BS performed experiments. MGK performed experiments, advised on study design, and edited the manuscript. PNT supervised the study.

## Acknowledgments

We would like to thank J. Wade Harper (Harvard Medical School) for generously gifting the RPS3 and RPL29 constructs for HALO editing HALO to the respective loci. We would also like to thank Ferhat Alkan and William J. Faller (Netherlands Cancer Institute) for their help with the ribosome heterogeneity analyses and their interpretation. Additionally, we wish to thank Claire L. Moore, Philip W. Hinds, and Karl Munger (Tufts University School of Medicine) for their feedback on the project as it was being developed, along with Dario Palmieri and Lara Rizzotto as part of the Gene Editing Shared Resource (The Ohio State University Comprehensive Cancer Center) for helpful discussions and support. Finally, we wish to thank Amanda E. Toland (The Ohio State University) for comments on the manuscript.

The work was supported by NCI grant P30 CA016058 grant to the Ohio State University, Comprehensive Cancer Center-OSUCCC. Additionally, MGK is supported by NIH grant R35GM146924.

