## Supplementary Materials and Methods for "KDM2B is required for ribosome biogenesis and its depletion unequally affects mRNA translation"

### Cell Culture

#### *Cells and culture conditions*

MDA-MB-231 cells (ATCC Cat: HTB-26) and HEK293T-LentiX cells (Takara Cat: 632180) were cultured in High Glucose Dulbecco's modified Eagle's minimal essential medium (Sigma Cat: D5796) supplemented with 10% fetal bovine serum (FBS) (Corning Cat: 35-010-CV), penicillin 100u/mL and streptomycin 0.1mg/mL (Sigma Cat: P4333), sodium pyruvate 1mM (Gibco Cat: 11360070), non-essential amino acids (Sigma Cat: M7145), and Plasmocin™ 5 µg/mL (Invivogen Cat: ant-mpp). Cells were subcultured following trypsinization (Trypsin-EDTA, Sigma Cat: T4049) every 2-3 days, depending on confluence and avoiding confluency >90%.

To validate cell line identity, genomic DNA extracted from individual lines with the DNEasy Blood and Tissue Kit (Qiagen Cat: 69504), was subjected to Short Tandem Repeat (STR) profiling (1) at The Ohio State University Comprehensive Cancer Center Genomics Shared Resource (Columbus, OH). The STR profiles of our cell lines matched more than 95% the profiles of the corresponding standard cell lines. This match was significantly higher than the 80% match, which is considered sufficient for cell line validation (2). Validation was repeated yearly to ensure the maintenance of cell line identity. Cell lines were also regularly tested for mycoplasma contamination, using a PCR-based assay (Mycoplasma Detection Kit, abm Cat: G238)

#### *Transient Transfection and Lentiviral Transduction*

Transfection of plasmids was performed using Lipofectamine 3000™, according manufacturer's protocol (Thermo Scientific Cat: L3000015). Briefly, 12 µg of DNA was added to 48 µl of P3000 to a final volume of 500 µl of . OptiMEM™ (Thermo Fisher Scientific Cat: 31985070), The P3000-diluted DNA was then added slowly to 500 µl of OptiMEM™ (Thermo Fisher Scientific Cat: 31985070), supplemented with 24 µl of Lipofectamine™. Following a 15-minute incubation at room temperature, the final mixture was added dropwise to the transfected cells. For cell lines with low transfection efficiency, we employed electroporation, using the SE Cell Line 4D X Kit (Lonza Cat: V4XC-1012) in a 4D Nucleofecter™ (Lonza Cat: AAF-1003X). Transfected DNA constructs, including gene editing constructs, are listed in Table 1.Lentiviral packaging was performed by transient, Lipofectamine-based transfection of lentiviral constructs and 2$^{nd}$ generation packaging plasmids (7.2µg of pPax2 (https://www.addgene.org/12260/) and 4.8µg of pMD2.G (https://www.addgene.org/12259/)) into 40-60% confluent Lenti-X cells in 10 cm dishes. 12-16 hours after transfection the media was removed from the plate and replaced with fresh media. Viruses were collected and filtered through 0.45µm filters (Corning Cat: 431220) at 48 and 72 hours from the start of the transfection. Harvested viruses were stored at -80 $^{0}$C. Cells were transduced with the harvested viruses, by overnight exposure to undiluted viral supernatants, supplemented with hexadimethrine bromide (polybrene) (Sigma Cat: 107689) to a final concentration of 8µg/mL. Selection of transduced cells for the appropriate antibiotic resistance was initiated 48 hours from the start of the exposure to the viruses. MDA-MB-231 cells transduced with constructs carrying the puromycin resistant gene were treated with 10µg/mL puromycin (Thermo Fisher Scientific Cat: A1113803) for 2-3 days. Cells transduced with constructs carrying the blastocidin resistance gene were treated with 10µg/mL of blastocidin (Thermo Fisher Scientific Cat: A1113903) for 5-7 days. Following selection, cells were cultured in antibiotic-free media.

**Immunoblotting**

Cells were lysed in Radioimmunoprecipitation Assay (RIPA) buffer (25mM Tris-HCl pH 7.6, 150 mM sodium chloride, 1% Nonidet P-40, 0.1% sodium deoxycholate, 0.1% SDS, 5mM EDTA), supplemented with Halt™ Protease and Phosphotase Inhibitor Cocktail (Thermo Fisher Scientific Cat: 78440), and lysates were collected by scraping with a cell lifter (Corning Cat: 3008). The protein lysates were vortexed, shaked for 30min at 4°C to complete the lysis and centrifuged at 15,000 g for 15 minutes at 4°C. Supernatants were collected in a fresh microcentrifuge tube. Protein concentration was determined, using the Pierce™ BCA Protein Assay Kit (Thermo Fisher Scientific, 23225). Lysates were then diluted 1:4 in 4X Laemmli Sample Buffer (BioRad Cat: 1610747) with 50 mM dithiothreitol (DTT). Following boiling for 10 minutes to complete protein denaturation, samples were loaded onto 4-15% gradient Mini-Protean TGX™ gel (BioRad Cat: 4561083/4561086), along with the protein ladder (BioRad Cat: 1610374). Gels were run in an electrophoresis chamber (BioRad Cat: 1658030) using SDS running buffer (25 mM Tris, 192 mM glycine, 0.1% SDS, pH 8.3) for 1-2 hours at a constant 100-140V.

Following electrophoresis, protein lysates were transferred to PVDF membranes (Thermo Fisher Scientific Cat: 88520) for 2-3 hours at 4°C, using a transfer buffer containing 25 mM Tris, 192 mM glycine, 20% v/v methanol and a constant current of 400 mA. For proteins larger than 200 kDa the transfer was carried out overnight at 4°C, at a constant current of 150 mA. After the transfer, membranes were stained with the Revert™ 700 Total Protein stain (Licor Biosciences Cat: 296-11020) and imaged on the 700 nm channel on a LI-COR Odyssey® Fc Imaging System (Licor Biosciences). Following this, membranes were washed in Tris-Buffered-Saline with Tween-20 (TBST) Buffer (20mM Tris-HCl, 150mM NaCl, 0.1% v/v Tween® 20), and blocked for 1 hour with 5% w/v BioRad Blocking Buffer (BioRad Cat:1706404), a milk-based blocking buffer in TBST, or with 5% w/v bovine serum albumin (BSA) (Fisher Scientific Cat: BP1600-100) in TBST. Membranes were then cut, based on the size of the proteins we planned to probe for, and the membrane strips were incubated at 4°C with the primary antibody in 5% w/v BSA overnight. The following day the primary antibodies were removed, and the membrane strips were washed 4-5 times in TBST, 5 minutes each time. Following incubation with horseradish peroxidase (HRP)-conjugated species-specific (rabbit or mouse) secondary antibodies in 5% w/v BSA for one hour at room temperature, membrane strips were washed 4-5 times in TBST (5 minutes each time) and incubated in either Pierce™ ECL2 Blotting Substrate (Thermo Fisher Scientific Cat: 80196) or SuperSignal™ West Femto Maximum Sensitivity Substrate (Thermo Fisher Scientific Cat: 34095). Membrane strips were finally imaged on a LI-COR Odyssey® Fc Imaging System, using the 700 nm channel for the protein ladder and the chemiluminescence HRP signal channel for test protein detection. Primary antibodies used for immunoblotting are listed in Table 2.

**RNA sequencing and data analysis**

RNA was extracted from 50-70% confluent MDA-MB-231 cells transduced either with a non-hairpin control shRNA or an shRNA targeting *KDM2B,* using the Total RNA Purification Plus Kit (Norgen Biotek Corp. Cat: 48300). Total RNA extracted was run on an Agilent Tapestation system with RNA Screentape™ devices (Agilent Technologies Cat: 5067-5576). RNAs with RNA Integrity Number (RIN) > 9.5 were submitted for library preparation.

RNA library preparation and sequencing were performed by the Brigham Young University DNA Sequencing Center (Provo, UT), via contract with Genohub Inc. (Austin, TX). Libraries were prepared using a KAPA RNA HyperPrep kit with RiboErase (Roche Cat:

KK8560) and they were subjected to paired-end sequencing (2x150bp) on an Illumina HiSeq Rapid V2 Chip, yielding an average of approximately 35 million reads per sample.

Reads were first analyzed for quality. Library preparation adapters were trimmed from the reads with TrimGalore (https://github.com/FelixKrueger/TrimGalore), which uses FASTQC (3, 4) to visualize sequencing quality and CutAdapt (5) to trim adaptor sequences. MultiQC was used to visualize sequencing quality across all samples (4).

The alignment of reads to the genome, gene quantification of reads, and differential analysis comparing control and *KDM2B* knockdown cells was done using RNAdetector (6). Reads were aligned to the *Homo sapiens* genome (GRCh38) using STAR (7) and aligned reads were then quantified using featureCounts (8). Differential expression analysis was performed in RNAdetector utilizing DESeq2 (9), edgeR (10), and LIMMA (11). The results of all 3 differential expression analysis tools were combined to generate a meta-p-value that was congruent across all 3 calculators using metaSeqR (12).

***Gene Set Enrichment Analysis***

Gene Set Enrichment Analysis (v4.3.2) (GSEA) (13, 14) was performed using the reads per million (RPM) table of the three control and three *KDM2B* knockdown replicate samples. The geometric mean of the control and shKDM2B samples was calculated for each gene, and genes with geometric means of $< 1$ were filtered out, as recommended by GSEA, to remove poorly expressed genes. This resulted in the retention of 13250 genes, corresponding to 12538 protein-encoding genes. Enrichment analysis was performed across the Gene Ontology (GO) Biological Processes dataset (15) that has gene lists of >15 and <500 genes in each category as the default setting for GSEA. 1000 permutations were run for each gene set, and enrichment scores were calculated for each permutation and compared to the enrichment score of the input-ranked list. Statistical significance was calculated based on the probability that the input ranked list had a better absolute enrichment score than the 1000 permutations ran. Statistical significance was adjusted for multiple hypotheses testing based on the total number of gene sets we considered.

***Exon Intron Split Analysis***

Using the Exon Intron Split Analysis (eisaR) software package (https://github.com/fmicompbio/eisaR) (16, 17), we calculated the fold change in read counts of the exons and introns of all genes in control and shKDM2B cells. Briefly, using the eisaR package and the human genome assembly (GRCh38) we generated genome annotation files that had labeled exons and introns across all protein-encoding gene bodies. We then aligned the RNAseq reads to the genome, using reads mapping to the exons along with reads mapping to the entire gene body. The reads mapping solely to the introns were determined by subtracting the reads mapping to the exons from the reads mapping to the entire gene body. Fold changes were calculated based on the exon or intron read counts between control cells and shKDM2B cells. These fold changes were then compared to each other to determine whether the fold changes in intron and exon read counts are concordant or discordant.

**Chromatin Immunoprecipitation and Sequencing (ChIP-Seq)**

Samples for Chromatin immunoprecipitation sequencing (ChIP-seq) experiments were prepared using the SimpleChIP Plus Enzymatic chromatin IP kit (Cell Signaling Technology Cat: 9005). 4 million MDA-MB-231 cells were cross-linked with 1% formaldehyde (Sigma-Aldrich Cat: F8775) for 10 min at room temperature. For the KDM2B ChIP we performed dual cross linking by treating the cells at room temperature for 20 minutes with 1.5 mM ethylene glycol-bis (succinimidyl succinate) (EGS) (Thermo Fisher Scientific Cat: 21565) dissolved in ice-cold PBS, and with 1% formaldehyde for 10 minutes, also at room temperature. Following chromatin digestion with 0.5 µL micrococcal nuclease, the nuclei were lysed by sonication with 3 cycles of 20s-on 30s-off pulses at 20% amplitude. 2% of the isolated chromatin was used prior to immunoprecipitation to extract control non-immunoprecipitated DNA.

ChIP-Seq libraries were prepared using the DNA Library Prep Kit (Cell Signaling Technology Cat: 56795) and Multiplex Oligos (Dual Index Primers) (Cell Signaling Technology Cat: 47538). PCR amplification and barcoding were carried out, using 5 ng of ChIP-enriched input DNA (for KDM2B), 1 ng input DNA (for c-MYC) and 50 ng input DNA (for H3K4me3, H3K27me3, H3K27ac and H2AK119ub). Finally, 50 ng of the DNA extracted from the non-immunoprecipitated chromatin was used as control. Antibodies used for ChIP, and the dilutions and volumes used, are listed in Table 3.Immunoprecipitated and control DNA libraries underwent paired-end sequencing (2x150bp), using an Illumina NovaSeq S4 Chip. DNA sequencing was carried out by Fulgent Genetics (Temple City, CA) via contract with Genohub Inc (Austin, TX) and yielded an average of 60 million reads per sample.

Library preparation adapters were first trimmed using CutAdapt (5), and reads were inspected for quality using FASTQC (3, 4). MultiQC was used to visualize sequencing quality across all samples (4). Reads were aligned to the *Homo sapiens* genome (GRCh38) using bowtie2 (18) and peak calling was performed using MACS2 (19). Visualization of global patterns of KDM2B and c-MYC binding, as well as global patterns of histone mark abundance, was done using the FLUFF tool (20).

**Assay for Transposase Accessible Chromatin Sequencing and Analysis**

$9x10^4$ MDA-MB-231cells were washed once with 1xPBS, and processed using the ATACseq kit (Active Motif, Cat: 53105). Briefly, cells were lysed with a lysis buffer that breaks down the plasma membrane but leaves the nuclei intact. Following centrifugation, the nuclei were incubated with a hyperactive mutant of the Tn5 Transposase and sequencing adapters. This results in DNA fragmentation and adapter ligation, selectively in open chromatin regions (tagmentation). Tagmented DNA was then purified and used for PCR amplification of the tagmented regions, with primers that were barcoded with different barcodes for each sample. To assess DNA concentration and amplicon size, the resulting libraries were purified with SPRI paramagnetic beads, which bind DNA fragment of a specific size range, selectively and reversibly and they were analyzed on an Agilent Bioanalyzer High Sensitivity DNA Chip (Agilent Technologies Cat: 5067-4626). Sequencing was performed by Fulgent Genetics (Temple City, CA) via contract with Genohub Inc (Austin, TX). ATAC-Seq libraries underwent paired-end sequencing (2x150bp), using an Illumina NovaSeq S4 Chip. DNA sequencing was carried out by Fulgent Genetics via contract with Genohub Inc and yielded an average of 240 million reads per sample. Sequencing data were analyzed, as described above for the ChIP-Seq data. Differential peak analysis between control and shKDM2B samples was performed using the DiffBind tool (21).

**Breast Cancer data analyses**

Gene expression in The Cancer Genome Atlas (TCGA) BRCA dataset (22), and the Genotype-Tissue Expression (GTEx) dataset (23) was obtained from the University of Santa Cruz (UCSC) Xena platform (24). RNA expression data from these datasets were normalized with the upper quartile method (25, 26). Protein expression data were from a subset of BRCAs in the TCGA database, whose proteome was analyzed by the Clinical Proteomic Tumor Analysis Consortium (CPTAC) using mass spectrometry (27). Correlations of the expression of KDM2B with the expression of genes encoding ribosome biogenesis factors at the RNA level were calculated from the TCGA transcriptomic data, and correlations at the protein level were calculated from CPTAC proteomic data. Scatter plots, Spearman correlation coefficients, and statistical significance were as reported in cBioportal (28, 29). BRCAs in the TCGA database that belong to different PAM50 subgroups had been identified in the original PAM50 paper (30). Correlations of the expression of KDM2B with the expression of genes encoding ribosome biogenesis factors were also calculated from the transcriptomic data of a large cohort of TNBCs (465 TNBCs, 360 with transcriptomic data), which was reported from the Fudan University Shanghai Cancer Center (FUSCC) (31). Fragment per kilobase million (FPKM) count matrices were downloaded from Bio-Med Big Data Center (www.biosino.org) under the node identification OEZ000398. The subclassification of these TNBCs into basal and non-basal was made by the group that published the original characterization of these tumors and was based on the tumor transcriptomic data.

***Pre-Ranked GSEA, based on correlations between KDM2B and the entire transcriptome of human BRCAs.***

Two sets of correlations between the *KDM2B* RNA levels and the RNA levels of all expressed genes were downloaded from the breast cancer database in cBioportal. The one set included all BRCAs, while the other one included only the basal-like BRCA subgroup (30). The same was done for the KDM2B protein levels and the levels of all the detected cellular proteins. However, the protein correlations were limited to the set of all the available BRCAs because the number of tumors in the basal-like subset was too small to generate accurately ranked correlation coefficients. The RNA gene expression and protein expression correlation coefficients with KDM2B were ranked from the most positively correlated to the most negatively correlated. This pre-ranked list was used as input for the GSEA pre-ranked enrichment analysis (13). Similarly to the RNA-seq enrichment analysis, 1000 permutations were run for each gene set, and an enrichment score was calculated and compared to the enrichment score of our input-ranked list. Statistical significance was calculated based on the probability that the input ranked list had a better absolute enrichment score than the 1000 permutations ran. Statistical significance was adjusted for multiple hypotheses testing based on the total number of gene sets we considered.

**TMT Proteomics**

1.5 million control and 2.5 million shKDM2B-transduced MDA-MB-231 cells were plated onto 10cm cell culture dishes, and they were harvested the next day. At the time of harvesting the confluency of all cultures was about 50-60%. Prior to harvesting, cells were washed 2x in 1xPBS. Harvested cell pellets were lysed in 500 µL proteomics lysis buffer (8M Urea, 50mM triethylamonium bicarbonate) containing Halt™ Protease and Phosphotase Inhibitor Cocktail (Thermo Fisher Scientific Cat: 78440). The samples were sonicated for 30 seconds at an amplitude of 25% on ice and then quantified using the Pierce™ BCA Protein Assay Kit (Thermo Fisher Scientific, 23225).

Samples were prepared for LC-MS/MS analysis at Bioinformatics Solutions Inc. (Waterloo, Ontario, Canada). Briefly, samples were reduced with 10mM DTT (Sigma-Aldrich, Missouri, USA), alkylated with 20 mM Iodoacetamide (Sigma-Aldrich, Missouri, USA) and precipitated in Acetone at -80°C. After removing the Acetone, the samples were then digested overnight with MS grade trypsin (Promega, Wisconsin, USA). Digested peptides were desalted with in-house made C18 spin columns and labeled with TMTpro 16plex (Thermo Fisher Scientific, Massachusetts, USA) following manufacturer's protocol. The labeled samples were quenched with Hydroxylamine and pooled to dry down. The pooled sample was then fractionated by high pH reverse phase into 44 fractions, and then combined into 11 samples.

Samples were resuspended in 12 µL buffer A (0.1% FA). For each run, 6 µL of each sample was separated by nanoflow liquid chromatography using an Ultimate 3000 chromatography system (ThermoFisher, Massachusetts, USA), then injected into the Thermo Orbitrap Fusion Lumos (ThermoFisher, Massachusetts, USA). Liquid chromatography was preformed using a constant flow of 0.25 µL/min and a 15 cm reversed-phased column with a 75 µm inner diameter filled with Reprosil C18 (PepSep, Bruker, Germany). Mobile phase A was 0.1% Formic Acid and Mobile phase B was 99.9% Acetonitrile, 0.1% Formic Acid. The separation was carried out over 120 minutes as follows: linearly 4% B to 35% B over 100 minutes with an increase to 95% B over 0.1 minute and held constant for 9.9 minutes to clean the column. Then the B percentage was set back to 4% in the final 10 minutes.

Two rounds of MS data were acquired on Thermo Orbitrap Lumos for each sample to maximize the proteome coverage. Both rounds were carried out in data-dependent mode with a cycle time of three seconds. In the first round, MS1 scan data were obtained at 120,000 resolution (at 400 m/z) with a mass range of 500–1,800 m/z. The automatic gain control (AGC) was set to standard, with an auto maximum ion injection time. The radio frequency (RF) lens was set to 30%. The charge state filter was set to 2-6 and the dynamic exclusion was set to 30 seconds. Isolation for MS2 scans was performed in the quadrupole, with an isolation window of 0.7 Da. MS2 scan data were acquired in the ion trap with CID activation at 30% collision energy (CE) and 10 millisecond (ms) activation time. The scan rate was set at Turbo with a standard AGC target and 35 ms ion injection time. The scan range of MS2 was set to auto and the mass range was set to normal. MS3 scans were performed in Orbitrap with HCD at 55% CE. The Orbitrap resolution was set to 50,000 and 10 synchronous precursor selections were used. AGC Target was set at 200% and maximum injection time was set at 200 ms. The parameters used in the second-round data acquisition were the same as the first-round, except the mass range in MS1 was set at 300-1,600 m/z and the CE was set at 45% for MS3 activation.

MS Raw Files were processed using PEAKS Studio XPro (Bioinformatics Solutions Inc., Ontario, Canada). The data was searched against the unreviewed human Unipot database. Precursor ion mass error tolerance was set to 15 ppm and fragment ion mass error tolerance was set to 0.6 Da. Quantification mass error tolerance (MS3) was set to 0.02 Da. Semi-specific cleavage with trypsin was selected with a maximum of 3 missed cleavages. A fixed modification of carbamidomethylation (+57.02 Da) on cysteine residues, and TMT16plex (+304.2072) on lysine and the peptide N-terminus were specified. Variable modifications of deamidation (+0.98 Da) on asparagine and glutamine, as well as oxidation (15.99 Da) on methionine were specified. A 1% false discovery rate (FDR) was set for the database search, and only proteins having at least 1 unique peptide and a fold change greater than 1 were reported.

**CRISPR Cas9 Gene Editing**

In frame fusion of the *RPS3* and *RPL29* endogenous loci with a HA LO-tag was performed, with sgRNA oligos cloned into pX330-U6-Chimeric_BB-CBh-hSpCas9-hGem vector (https://www.addgene.org/71707/) and targeting the 3' ends of these loci, proximal to the stop codon (32-34). Cas9 was expressed from this vector as a fusion with a geminin tag, to ensure that Cas9 would only be expressed in the S or G2 phases of the cell-cycle, after the DNA has been replicated, increasing the chance for homology director repair (35). The donor plasmids contained the HALO-tag, which was inserted between the two *RPS3,* or *RPL29* homology arms, downstream of a flexible serine-glycine rich linker. These plasmids were a generous gift from J.

Wade Harper (32). Constructs used for transfection of Cas9 and the sgRNA are listed in Table 1 and of sgRNA design are listed in Table 4.

The sgRNA and Cas9-containing plasmid and the donor plasmids were co-transfected into MDA-MB-231 cells, using Lipofectamine 3000™. Following transfection, and after they were given sufficient time to divide, cells were treated with the fluorescent Halo-Ligands JFX-549 or JFX-650 (Janelia Research Campus) (36, 37). Fluorescent cells were sorted on a SH800S Cell Sorter (Sony Cat: SH800S) operated by the Gene Editing Shared Resource of The Ohio State University Comprehensive Cancer Center (Columbus, OH). Single cell clones were cultured in 96-well plates (Corning, Cat: 3596). Individual clones were analyzed by western blotting to confirm proper editing.

*RPS3-Halo* and *RPL29-Halo-e*dited clones were transduced with pLKO-shKDM2B or the empty vector. Following puromycin selection, the transduced cells were plated onto 60mm dishes. One replicate plate of shKDM2B and EV-transduced cells was used to count the number of cells per plate at the start of the experiment. Based on the numbers of transduced MDA-MB-231 cells and the published protein molarity data per cell in OpenCell (38), we estimated the molar amounts of RPS3 and RPL29. Following this, cells were stained to saturation with JFX-549, used at a molar concentration 10x higher than the molar concentration of RPS3, or RPL29. After 1 hour of exposure to JFX-549, the cells were washed twice with room temperature 1xPBS and then pulsed for 24 hours with the halo fluorescent ligand JFX-650, which labeled newly assembled ribosomes (32). Given that shKDM2B-transduced MDA-MB-231 cells grow slowly (39), the chase of the JFX-650 pulse was extended to more than 24 hours, to reliably capture the rate of assembly of new ribosomes.

**Global mRNA translation using Biorthogonal Amino Acid Labeling of newly synthesized proteins.**

Cells grown to 60% confluence in 10cm dishes were starved of L-methionine, L-cystine, and L-glutamine for 1 hour by culturing them in DMEM lacking all three amino acids (Sigma Cat: D0422) and supplemented with 10% dialyzed FBS (40). After starvation, the media was replaced with DMEM containing L-cystine, L-glutamine, and 100μM L-homopropargylglycine (L-HPG) (Click Chemistry Tools Cat: 1067) and supplemented with 10% FBS and they were harvested at the indicated time points, by scrapping in 1mL of a lysis buffer containing 0.16% SDS in 1xPBS with Halt™ Protease and Phosphatase Inhibitor Cocktail (Thermo Fisher Scientific Cat: 78440). Protein concentration was determined using the Pierce™ BCA Protein Assay Kit (Thermo Fisher Scientific, 23225). Two mg of protein of each sample were then loaded into a fresh tube and 1x PBS was added to a final volume of 2 mLs. Newly synthesized, L-HPG-labeled proteins in these samples were detected, using click chemistry.

Click-chemistry solutions were prepared as 100x stocks and they included: Biotin-Azide, (10mM PEG4 carboxamide-6-Azidohexanyl Biotin), (Click Chemistry Tools Cat: 1265), TBTA (10mM Tris[(1-benzyl-1H-1,2,3-triazol-4-yl) methyl]amine) (Sigma Cat: 678937), TCEP (100mM Tris (2-carboxyethyl) phosphine) (Sigma Cat: C4706), and 100mM Copper (II) Sulfate (Sigma Cat: C1297). TCEP and Copper (II) Sulfate stocks were made fresh the day of the click-chemistry reaction while Biotin-Azide and TBTA were stored in -80 $^{o}$C. 20μL of each of the stock solution were added to each of the 2mL L-HPG-labeled samples and they were mixed with the samples by shaking for 2 hours in the dark at room temperature. Following this, 10 mL of pre-chilled acetone (-20 $^{o}$C) was added, and samples were placed in -20$^{o}$C for the proteins to precipitate overnight. The next day samples were centrifuged at 4000xg for 30min at 4$^{o}$C, the acetone was removed, and the precipitates were air-dried and resuspended in 700 μL of a 1.2% SDS solution, containing Halt™ Protease and Phosphatase Inhibitor Cocktail (40). Samples were then electrophoresed and electroblotted onto PVDF membranes. The membranes were stained using Revert™ 700 Total Protein stain (LicorBiosciences Cat: 296-11020), and they were probed with an anti-biotin antibody to detect biotinylated nascent peptides.

**Polysome Profiling**

Control and shKDM2B-transduced MDA-MB-231 cells, were harvested at 16 hours after plating, from cultures growing at similar densities. Given that the rate of growth of shKDM2B cells lags behind the rate of growth of control cells (41), we fist addressed the relative numbers of the two cell types we had to plate, to produce equal numbers of cells at the time of harvesting, and we observed that this is accomplished by plating 2.7 million control cells and 4.5 million shKDM2B-transduced cells into 10 cm dishes. At the time of harvesting, cells plated 16 hours earlier were placed on ice and they were washed twice with ice-cold 1xPhosphate buffered saline (PBS), (Sigma Cat:D8537) containing Cycloheximide (CHX) (Sigma Cat:C4589) (100μg/m). After the last wash, the cells were scraped gently into 500μL of PBS, and they were centrifuged at 300xg, at 4$^{o}$C for 5 minutes. Liquid supernatants were then removed, and cell pellets, containing approximately 40-50 million cells, were flash-frozen by immersion into liquid nitrogen and stored at -80 $^{o}$C. Three biological replicates were generated and stored for each condition.

Polysome profiling was performed, by following a standard protocol (42). A 50% sucrose solution (50% w/v sucrose,10mM Tris pH 7.4, 140mM KCl, 10mM $MgCl_2$, 1mM DTT, and 100μg/mL CHX) was placed in the bottom of an SW41 ultracentrifuge tube and it was carefully overlayed with a 10% sucrose solution (10% w/v sucrose, 10mM Tris pH 7.4, 140mM KCl,

10mM $MgCl_2$, 1mM DTT, and 100μg/mL CHX). Using a Gradient Master (Biocomp Cat: 108), the two solutions were mixed to generate a continuous 10-50% sucrose gradient. The frozen cell pellets were lysed in polysome lysis buffer (1% Triton-X100v/v 20mM Tris pH 7.4, 140mM KCl, 10mM $MgCl_2$, 1mM DTT, and 100μg/mL CHX). The volume of the lysis buffer was adjusted to the estimated number of cells in the pellet, with the lowest volume being 400 μL. Cell lysates were mixed, vortexed, and placed on ice for 10min. Following this, they were centrifuged at 2000xg and the supernatant was collected. 400μL of the supernatant was carefully loaded onto the sucrose gradient and ultracentrifuged at 260,000xg/x $^0$C for 2 hours with slow acceleration and deceleration (no brakes). Gradient fractions were collected from top to bottom (lightest to heaviest), using a gradient fractionator (Biocomp Cat: 152), with absorbance measured on at 260 nm, with a Triax 3 wavelength flow cell (Biocomp Cat: FC-3-260-2VIS).

**Paired RNA sequencing and Ribosome Protected Fragment sequencing and analysis.**

***Sample preparation and RNA sequencing***

1.5 million control and 2.5 million shKDM2B-transduced MDA-MB-231 cells were plated into 10 cm cell culture dishes to achieve confluence of 50-60% for both at the time of harvesting 16 hours later. Prior to harvesting, cells were placed on ice and they were washed twice with ice-cold 1xPhosphate buffered saline (PBS), (Sigma Cat:D8537) supplemented with 100μg/mL Cycloheximide (CHX) (Sigma Cat:C4589). Following the last wash cells were scraped gently into 500 μL of PBS and they were collected and centrifuged at 300xg for 5 minutes at 4$^o$C. Supernatants were then quickly pipetted off and the cell pellets were flash-frozen by immersion in liquid nitrogen and stored at -80 $^o$C until ready for polysome profiling. Each cell pellet contained 10-12 million cells. Three biological replicates were collected for each experimental condition.

Library preparation and sequencing were performed by TB-Seq™ (San Francisco, CA). Briefly, cells were resuspended in ice cold lysis buffer (20 mM TrisHCl, pH 7.4, 100 mM NaCl, 5 mM $MgCl_2$ 1% v/v Triton X 100, 1 mM DTT, 20U/mL Turbo DNase I, 0.1 % NP40, 100 μg/mL cycloheximide) and the soluble cytoplasmic fractions were collected following centrifugation for 20 min at 4 °C at top speed in a microcentrifuge (43, 44). Clarified cytoplasmic fractions were then digested with RNase I for 45 min at room temperature. Digestion was stopped with SUPERase-IN™ (Invitrogen Cat: AM2694) and monosomes were purified by size exclusion chromatography on MicroSpin S-400 HR columns (GE Healthcare Cat: GE27-5140-01) as described (45). Size selection of 25-36 nt footprints was performed by electrophoresis in 15% TBE-urea gels. Illumina ready RIBO-seq libraries were prepared using a SMARTer smRNA-seq kit (TakaraBio Cat: 645031). Library cDNA concentrations were measured by qubit fluorometer, and the quality of the libraries was assessed on a Agilent 2100 bioanalyzer. RIBO-seq libraries were sequenced on an Illumina Novaseq 6000 sequencer, producing an average of 90 million single-end reads per sample, 1x50 cycles.

RIBO-Seq, was paired to RNA-Seq, which was carried out in parallel with the RIBO-Seq, also by TB-Seq™ (San Francisco, CA). RNAs for RNA-Seq were isolated from aliquots of the same cell lysates, used for the RIBO-Seq experiment, and libraries were prepared after the total RNAs were depleted of rRNA, using a RIBO-Minus transcriptome isolation kit (Invitrogen Cat: A1083708). mRNA fragmentation aiming to generate RNA fragments of sizes similar to those of the ribosome footprints was carried out by incubating RNAs for 25 min at 94 °C. SMARTer smRNA-seq kit (TakaraBio Cat: 645031) was used to generate Illumina ready RNA-seq libraries. The assessment of the DNA concentration and quality of the constructed libraries, as well as the sequencing of the libraries were carried out as described for the RIBO-Seq libraries for an average of 50 million single-end reads per sample, 1x50 cycles

***The canonical transcriptome of MDA-MB-231 cells***

The short reads of the paired RIBO-Seq/RNA-Seq are more likely to map to multiple sites of the genome confounding the results and limiting the analysis (46). Additionally, transcription and RNA processing of most genes give rise to multiple RNA transcripts, and the short reads do not allow one to distinguish between different transcripts in a given cell. This information is important for the analysis of the RIBO-Seq data because the counting of ribosome-protected fragments should be limited to transcripts representing the "canonical" transcriptome of the cell (47, 48). Definitions vary on what defines a "canonical" transcriptome with some groups defining it based on the transcripts with the longest coding regions of each gene (47) and others defining it based on the most abundant transcripts of each gene (48). Here we adopted the latter definition, and we used an RNA-Seq with longer reads (2x 150 bps) we had already performed on control and shKDM2B-transduced MDA-MB-231 cells(49), to identify the most abundant transcripts in these cells. To do this, we used RNAdetector (6) to run SALMON (50) and obtain read counts at the transcript isoform level.

***Data Analysis***

We first used TrimGalore (https://github.com/FelixKrueger/TrimGalore) which employs FASTQC (3, 4) to visualize sequencing quality and CutAdapt (5) to trim adaptor sequences. Additionally, we used MultiQC to visualize sequencing quality across all samples (4). Trimmed reads were then aligned to the rRNA and tRNA transcriptome to remove these abundant transcripts from the RIBO-Seq datasets by using bowtie2. Reads that were unmapped to the rRNA and tRNA transcriptome were then aligned to the *Homo sapiens* genome (GRCh38) using

STAR (7) in both genome and transcriptome mode. The alignment in transcriptome mode was then analyzed using riboWaltz to determine the p-site offsets, which define the optimal p-site for each fragment length (51), and to address whether the p-site offsets displayed proper trinucleotide periodicity. RNA-seq and Ribo-seq mapped reads in genome mode were then quantified using featureCounts based on the identified canonical transcriptome(8). The paired RNA-seq and Ribo-seq reads were then analyzed by deltaTE (52) which uses DEseq2 (9) to calculate changes in the abundance of ribosome protected RNA fragments and changes in transcript abundance in the paired RNA-Seq. Normalization of the abundance of ribosome protected RNA fragments, based on the abundance of RNA transcripts, defined translational efficiency (TE). Changes in TE in shKDM2B cells (ΔTE) defined the relative sensitivity of the translation of individual transcripts to the loss of KDM2B. Gene ontology analysis of transcripts with altered translational efficiency were analyzed by Metascape (53).

Given that transcripts varied in ΔTE, we examined whether differences in ΔTE correlate with structural differences between transcripts. First, we examined how the length of the most abundant transcripts and the length and structure of their 5' UTRs correlate with their ΔTE. Transcript length and the sequence and structure of the 5' untranslated region (UTR) were downloaded from University of California Santa Cruz (UCSC) Genome Browser in transcript information and foldUTR5 databases, respectively (48). Next, we examined how known and de novo RNA motifs in the 5' and 3' UTRs affect the ΔTE of individual transcripts. Motif enrichment analysis was performed on meme-suite.org using the 5'UTR sequences of transcripts detected by the RIBOseq. Motif enrichment was determined by comparing the sequences of the translationally upregulated transcripts or downregulated transcripts to the transcripts that were not significantly altered translationally. Known RNA-binding motifs using the Human Ray2013

RNA binding protein motif database using Analysis of Motif Enrichment (AME) and de novo motifs were identified using Sensitive, Thorough, Rapid, Enriched Motif Elicitation (STREME).

RiboWaltz produces p-site density data (51), which can be used to identify potential differences in codon usage between two conditions. We therefore compared the p-site densities in all the control and all the shKDM2B biological replicates to address whether the loss of KDM2B alters the preference of codon utilization.

Selective changes in mRNA translational efficiency, like the ones observed in KDM2B knockdown cells, may be due to ribosomal heterogeneity (54), which can potentially be addressed by reanalyzing the RIBO-Seq data using dripARF (55). Based on the 3D structure of the ribosomes, the ribosomal RNA sequences neighboring different ribosomal proteins are known. Using this information, we can translate the abundance of ribosomal RNA fragments protected from RNAse I digestion because of their binding to ribosomal proteins into the abundance of such proteins in the ribosomes. We therefore reanalyzed our RIBO-Seq data, focusing on changes in the abundance of rRNA fragments, which are normally removed during RIBO-Seq data analysis.

**Northern Blotting**

***RNA extraction, electrophoresis, and transfer.***

RNA was extracted using Purelink™ RNA Mini kit (Invitrogen Cat: 12183018A) and it was analyzed for quality and quantity with an Agilent Bioanalyzer, using an RNA 6000 Nano Kit™ (Agilent Technologies Cat: 5067-1511). Northern blotting was performed using the NorthernMax™ Kit (Invitrogen Cat: AM1940) and following the manufacturer's protocol. Briefly, prior to loading, 1μg of total RNA was mixed with 3 volumes of formaldehyde loading dye and ethidium bromide to a final concentration of 10μg/mL and denatured at 65°C for 15 minutes.. RNA samples were loaded onto denaturing agarose gels made using the formaldehyde-based 10X denaturing gel buffer as per manufacturers protocol and RNAseq free water. The gel with RNA were electrophoresed in 1xMOPS buffer at 100V for 2 hours. Gels were imaged under ultraviolet light. Following electrophoresis, RNA was transferred to BrightStar™-Plus Positively Charged Nylon Membranes (Invitrogen Cat: AM10100) by capillary action, using the standard protocol provided by the manufacturer. The RNA was cross-linked to the membranes, using ultraviolet light emitted by a UV Stratalinker™ 2400 (Agilent Cat: Discontinued). RNA-loaded membranes were pre-hybridized in 10mL of ULTRAhyb™ Ultrasensitive hybridization buffer at 68°C in a heat-sealable bag for 1 hour.

***Probe synthesis, hybridization, washing and development.***

Four sets of primers were designed to amplify ~200nt sections of the two external and two internal transcribed spacers (3' and 5'ETS and ITS1/ITS2 respectively) of the rRNA genomic loci (56). The forward primer of the 3' ETS included 20 nucleotides from the 3'end of the 28S rRNA and as a result, the probe synthesized using this primer, also detected the mature 28S rRNA. Genomic DNA (gDNA) was amplified by PCR, using these primers and Phusion polymerase (New England Biolabs Cat: M0530S) or Platinum™ SuperFi II polymerase (Thermo Fisher Cat: 12361010). PCR products were electrophoresed in 1% agarose gels and they were extracted using the Nucleospin® PCR and gel clean-up kit (Macherey Nagel Cat: 740609). The extracted amplicon was reamplified, using the same forward primer, and a reverse primer with an overhang that added a T7 viral RNA polymerase promoter on its 3'end. Following gel electrophoresis and gel extraction as described above, amplified DNA was quantified using a NanoDrop™ One/OneC Microvolume UV-Vis Spectrophotometer. 200ng of the T7 promoter-containing PCR amplicon was used as the input for T7 polymerase-directed in vitro transcription, using the Digoxigenin (DIG) RNA Labeling Kit (SP6/T7) (Roche Cat: 11175025910). 10% of the product was electrophoresed in an agarose gel to determine the efficiency of the transcription reaction, and the remainder was stored in -80°C until ready to use.

The DIG-UTP labeled antisense probe was diluted in 1 mL hybridization buffer and was added to the hybridization bag. Hybridization was carried out at 68°C overnight. Hybridized membranes were washed twice, 5 minutes each time, in 2x SSC with 0.1% SDS (300mM NaCl, 30mM sodium citrate pH 7.0, 0.1% w/v SDS) at room temperature. Next, they were washed twice, 15 minutes each time, in 0.1XSSC with 0.1% SDS (15mM NaCl, 1.5mM sodium citrate pH 7.0, 0.1% w/v SDS) at 68°C, and following this, they were prepared for the detection of the hybridized probe with an enzyme-linked immunoassay, using the DIG-Starter Kit (Roche Cat:12-039-672-910). Briefly, membranes were washed twice, 5 minutes each time, with wash buffer (0.1 M maleic acid pH 7.5, 0.15 M NaCl, 0.3% v/v Tween Tween® 20) at room temperature, and then, they were incubated, with the blocking solution (0.1 M maleic acid, 0.15 M NaCl), for 30 minutes, also at room temperature. Subsequently, membranes were incubated for 1 hour at room temperatures, with an alkaline phosphatase-conjugated anti-digoxigenin antibody diluted 1:15,000 in blocking solution. Following two room temperature, washings, 15 minutes each, in wash buffer, and a final 5 minute wash in detection buffer (0.1 M Tris-HCl, 0.1 M NaCl, pH 9.5), membranes were placed on a plastic sheet, treated dropwise with the chemiluminescence substrate CDP-Star® (Roche Cat: 12041677001) and then covered with a plastic cover. Five to ten minutes later, membranes were imaged on the chemiluminescence channel on a LI-COR Odyssey® Fc Imaging System to capture the probe-antibody signal and the 600nm channel to capture the ethidium bromide signal. Exposure on both channels was for 10 minutes.

**Cell cycle distribution: Propidium Iodide staining and Flow Cytometry**

Cells were trypsinized, centrifuged at 300xg, and resuspended in 800 µl of 1x PBS. Following this, 2.5 mL of cold (-20 $^{0}$C) 100% ethanol was added dropwise to the cell suspension while vortexing at low speed. The ethanol/PBS-resuspended cells were kept at -20°C overnight. The next day, the cells were centrifuged at 300xg, and the cell pellets were washed once in 1x PBS and resuspended in 0.5 mL of FxCycle™ PI/RNase Staining Solution (Invitrogen Cat: F10797). Following a 30-minute incubation, we used flow cytometry (BD LSRFortessa™ Cell Analyzer), to determine the distribution of cells with different intensities of PI fluorescence, and the data were analyzed on FlowJo (v10) (https://www.flowjo.com/).

**Statistical software and analyses**

Statistical analyses and data plotting were performed using the GraphPad Prism Software (v9.5.1) (https://www.graphpad.com/), or RStudio (v2023.03.0+386) (https://posit.co/download/rstudio-desktop/). The statistical software and the statistical tests applied to the results of individual experiments are described in the figure legends.

Table 1

| Construct | Source | Usage |
|---|---|---|
| pLKO.1-Control | https://www.addgene.org/10879/ | Control plasmid with non-targeting shRNA |
| pLKO.1-shKDM2B #1 | Millipore Sigma pre-designed shRNA TRCN0000238779 | shRNA lentiviral plasmid targeting *KDM2B* |
| pLKO.1-shKDM2B #2 | Millipore Sigma pre-designed shRNA TRCN0000118437 | shRNA lentiviral plasmid targeting *KDM2B* |
| pLKO.1- shEZH2 | Millipore Sigma pre-designed shRNA TRCN0000018365 | shRNA lentiviral plasmid targeting *EZH2* |
| pLKO.1-shPCGF1 | Millipore Sigma pre-designed shRNA TRCN0000073128 | shRNA lentiviral plasmid targeting *PCGF1* |
| pLKO.1-shPCGF4 | Millipore Sigma pre-designed shRNA TRCN0000020157 | shRNA lentiviral plasmid targeting *PCGF4* |

| | | |
|---|---|---|
| **pLKO.1-** shEIF4G2 | Millipore Sigma pre-designed shRNA TRCN0000147956 | shRNA lentiviral plasmid targeting *EIF4G2* |
| pPAX2 | https://www.addgene.org/12260/ | 2nd generation lentiviral packaging plasmid |
| pMD2.G | https://www.addgene.org/12259/ | 2nd generation lentiviral packaging plasmid |
| pSMART RPS3-HALO-FLAG Homology Cloning Vector | Gift from J. Wade Harper (32) | Donor plasmid for the editing of RPS3 endogenous locus via CRISPR Cas9 |
| pSMART RPL29-HALO Homology Cloning Vector | Gift from J. Wade Harper (32) | Donor plasmid for the editing of RPL29 endogenous locus via CRISPR Cas9 |
| pSpCas9-2A-Puro | https://www.addgene.org/62988/ | Vector backbone to clone sgRNA into a transient puromycin selectable vector for knockouts or testing gRNA efficiency |
| pSpCas9-hGem | https://www.addgene.org/71707/ | Vector backbone to clone sgRNA into a transient transfection vector where Cas9 is fused to the degradation tag of Geminin |

Table 2

| **Antibody** | **Catalog Number** | **Concentration used** |
|---|---|---|
| anti-Biotin-HRP | Cell Signaling Technology Cat: 7075S | 1:2000 |
| anti-CCNB1 | Cell Signaling Technology Cat: 12231S | 1:1000 |
| anti-cdc2 (CDK1) | Cell Signaling Technology Cat: 9116S | 1:1000 |
| anti-EIF2α | Cell Signaling Technology Cat: 9722S | 1:1000 |
| anti-EZH2 | Cell Signaling Technology Cat: 5246S | 1:1000 |
| anti-FBL | Cell Signaling Technology Cat: 5246S | 1:1000 |
| anti-JHDM1B (KDM2B) | Millipore Cat: 09-864 | 1:1500 |
| anti-NAT10 | Proteintech Cat: 13365-1-AP | 1:1000 |

| | | |
|---|---|---|
| anti-Phospho-EIF2α Ser51 | Cell Signaling Technology Cat: 9721S | 1:1000 |
| anti-RPL29 | Proteintech Cat: 15799-1-AP | 1:1000 |
| anti-RPS28 | Thermo Scientific Cat: PA5-45721 | 1:1000 |
| anti-RPS3 | Cell Signaling Technology Cat: 9538S | 1:1000 |
| anti-TCOF1 | Proteintech Cat: 11003-1-AP | 1:1000 |
| anti-UTP14A | Proteintech Cat: 11474-1-AP | 1:1000 |
| anti-MYC | Cell Signaling Technology Cat: 18583S | 1:1000 |
| anti-PCGF1 | Thermo Scientific Cat: PA5-49390 | 1:1000 |
| anti-PCGF4 | Cell Signaling Technology Cat: 6964S | 1:1000 |
| anti-PKR | Cell Signaling Technology Cat: 3072S | 1:1000 |
| anti-phospho-PKR Thr446 | Millipore Cat: 07-532 | 1:1000 |
| anti-PERK | Cell Signaling Technology Cat: 3192S | 1:1000 |
| anti-EIF4G2 | Cell Signaling Technology Cat: 5169S | 1:1000 |
| anti-ETS1 | Cell Signaling Technology Cat: 14069S | 1:1000 |
| anti-IFIT5 | Cell Signaling Technology Cat: 85345S | 1:1000 |
| anti-CTNNB1 | Cell Signaling Technology Cat: 8480S | 1:1000 |
| anti-MMP1 | Cell Signaling Technology Cat: 54376S | 1:1000 |
| anti-c-JUN | Cell Signaling Technology Cat: 9165S | 1:1000 |
| anti-Vinculin | Cell Signaling Technology Cat: 13901S | 1:1000 |
| anti-H3 | Cell Signaling Technology Cat: 4499S | 1:3000 |
| anti-α-Actinin | Cell Signaling Technology Cat: 6487S | 1:2000 |
| anti-α-Tubulin | Sigma Cat: T5168 | 1:3000 |
| anti-β-actin | Cell Signaling Technology Cat: 4970S | 1:3000 |

Table 3

| Antibody | Catalog Number | Concentration used |
|---|---|---|
| anti-JHDM1B ChIP+ (KDM2B) | Millipore Cat: 17-10264 | 1:100 |
| anti-Myc | Cell Signaling Technology Cat: 18583S | 1:100 |
| anti-H3K4me3 | Cell Signaling Technology Cat: 9751S | 1:50 |
| anti-H3K27me3 | Cell Signaling Technology Cat: 9733S | 1:50 |
| anti-H2AK119ub | Cell Signaling Technology Cat: 8240S | 1:100 |
| anti-H3K27ac | Cell Signaling Technology Cat: 8173S | 1:50 |

Table 4

| Oligonucleotides | Name | Oligonucleotide sequence (5' to 3') |
|---|---|---|
| 5' ETS Cloning and T7 Addition from gDNA | 5' ETS Forward Primer | CCCGTGGTCTCTCGTCTTCT |
| | 5' ETS Reverse Primer | CTGTCCGGGAGGGACCAC |
| | 5' ETS Reverse Primer T7 addition | TAATACGACTCACTATAGGGctgtccgggagggaccac* |
| ITS1 Cloning and T7 Addition from gDNA | ITS1 Forward Primer | GCGAGAGCCGAGAACTC |
| | ITS1 Reverse Primer | GAGGAGGGCACCGAGACC |
| | ITS1 Reverse Primer T7 addition | TAATACGACTCACTATAGGGgaggagggcaccgagacc* |
| ITS2 Cloning and T7 Addition from gDNA | ITS2 Forward Primer | GTCCCCCTAAGCGCAGAC |
| | ITS2 Reverse Primer | GGCTCTCTCTTTCCCTCTCC |
| | ITS2 Reverse Primer T7 addition | TAATACGACTCACTATAGGGggctctctctttccctctcc* |
| 3' ETS Cloning and T7 Addition from gDNA | 3' ETS Forward Primer | CTCGACACAAGGGTTTGTCC |
| | 3' ETS Reverse Primer | AGGCGGGAACCGAAGAAG |

| | | |
|---|---|---|
| | 3' ETS Reverse Primer T7 addition | TAATACGACTCACTATAGGGaggcgg gaaccgaagaag* |
| RPS3 sgRNA Oligos | Strand Targeting Oligo | caccGACATACCTGTTATGCTGTG** |
| | Complementary Oligo | aaacCACAGCATAACAGGTATGTC** |
| RPL29 sgRNA Oligos | Strand Targeting Oligo | caccGAGATATCTCTGCCAACATG** |
| | Complementary Oligo | aaacCATGTTGGCAGAGATATCTC** |
