## Supplementary Figures and Legends for "KDM2B is required for ribosome biogenesis and its depletion unequally affects mRNA translation"

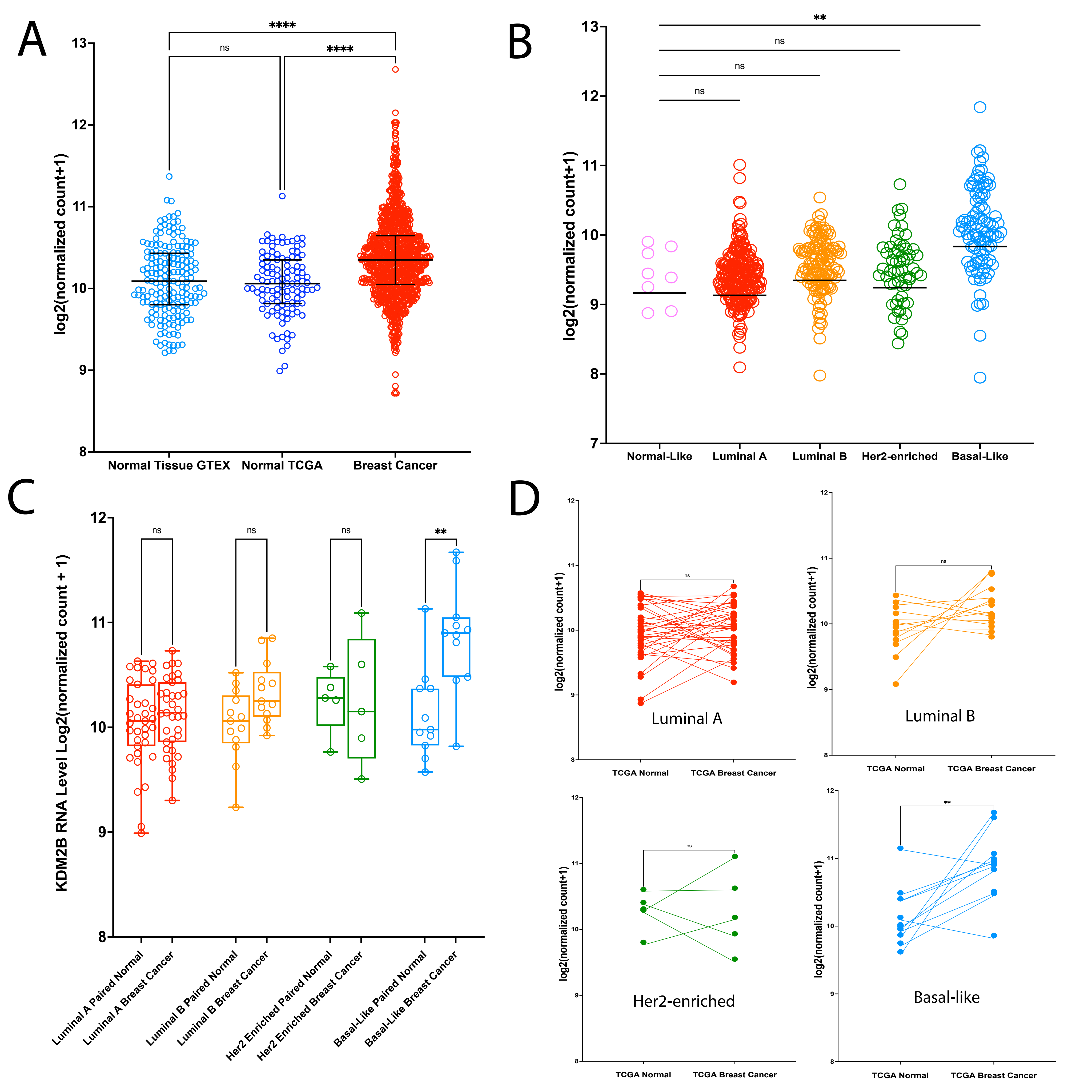
A
B
C
D
log2(normalized count+1)
Normal Tissue GTEX
Normal TCGA
Breast Cancer
ns
****
**
Normal-Like
Luminal A
Luminal B
Her2-enriched
Basal-Like
KDM2B RNA Level Log2(normalized count + 1)
Luminal A Paired Normal
Luminal A Breast Cancer
Luminal B Paired Normal
Luminal B Breast Cancer
Her2 Enriched Paired Normal
Her2 Enriched Breast Cancer
Basal-Like Paired Normal
Basal-Like Breast Cancer
TCGA Normal
TCGA Breast Cancer
Luminal A
Luminal B
Her2-enriched
Basal-like

**Supplementary Figure 1.** KDM2B is overexpressed in basal-like breast cancer

**A.** Comparison of KDM2B RNA levels between normal breast (GTEx (1) and TCGA (2)) and breast cancer. **B.** Comparison of KDM2B RNA levels in PAM50 breast cancer subtypes. For both A and B, statistical significance was calculated by Kruskal-Wallis, followed by Dunn's multiple comparisons test. **C.** KDM2B RNA levels in paired tumor and normal samples across the four PAM50 subtypes. **D.** Bipartite graphs of KDM2B levels in paired samples of tumor and normal breast. Wilcoxon signed-rank test was employed for the paired analyses (C and D). **p-value < 0.01, ****p-value <0.0001.

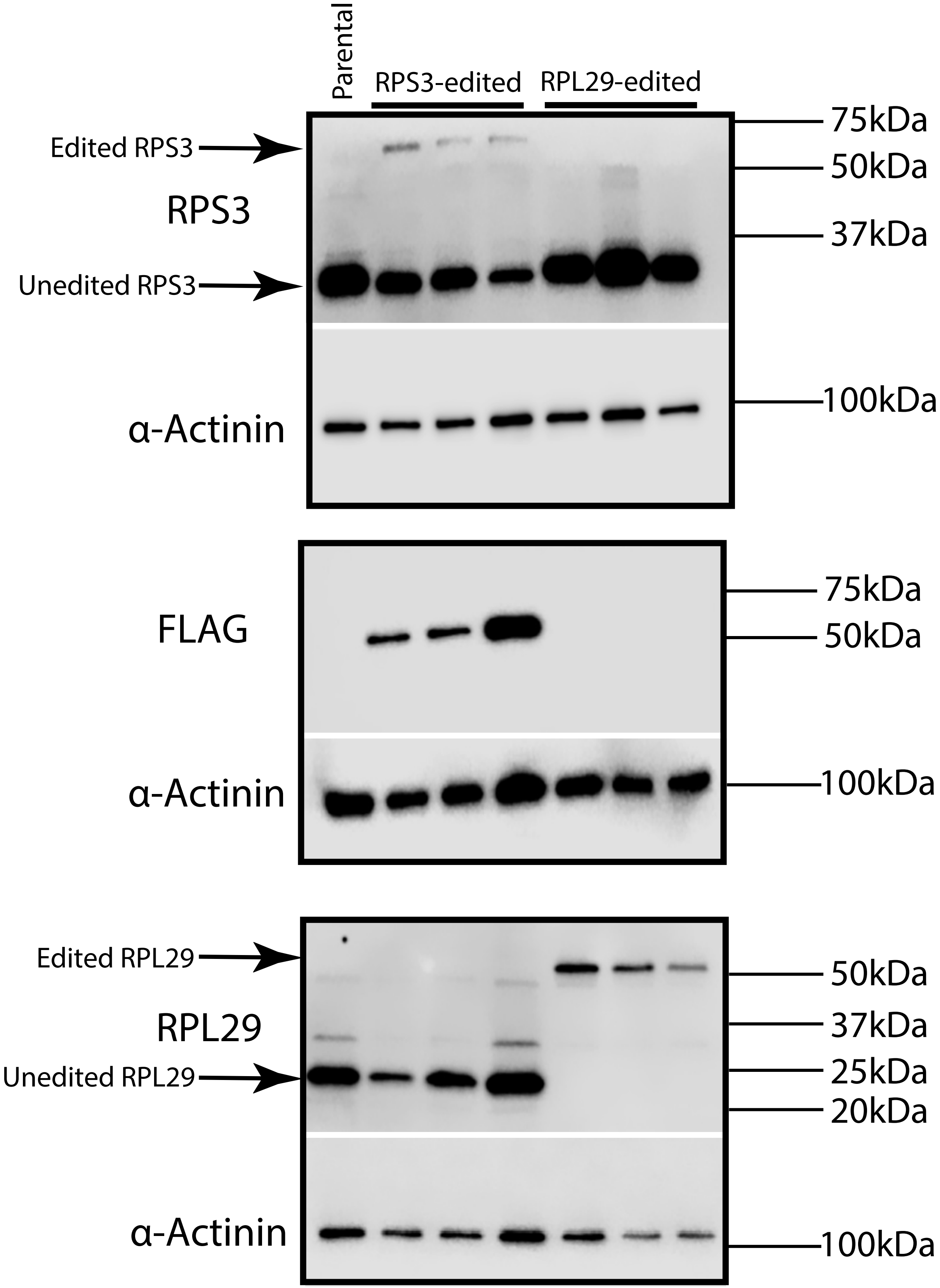
Parental
RPS3-edited
RPL29-edited
Edited RPS3
Unedited RPS3
RPS3
α-Actinin
75kDa
50kDa
37kDa
100kDa
FLAG
α-Actinin
75kDa
50kDa
100kDa
Edited RPL29
Unedited RPL29
RPL29
α-Actinin
50kDa
37kDa
25kDa
20kDa
100kDa

**Supplementary Figure 2.** Confirmation of Editing of the *RPS3* and *RPL29* loci.

Western blots of CRISPR-edited cells were probed with antibodies for RPS3 (Upper panel), FLAG (Middle panel), or RPL29 (Lower panel). RPS3 homology design also contained a FLAG tag, allowing for the detection of proper editing with the anti-FLAG antibody.

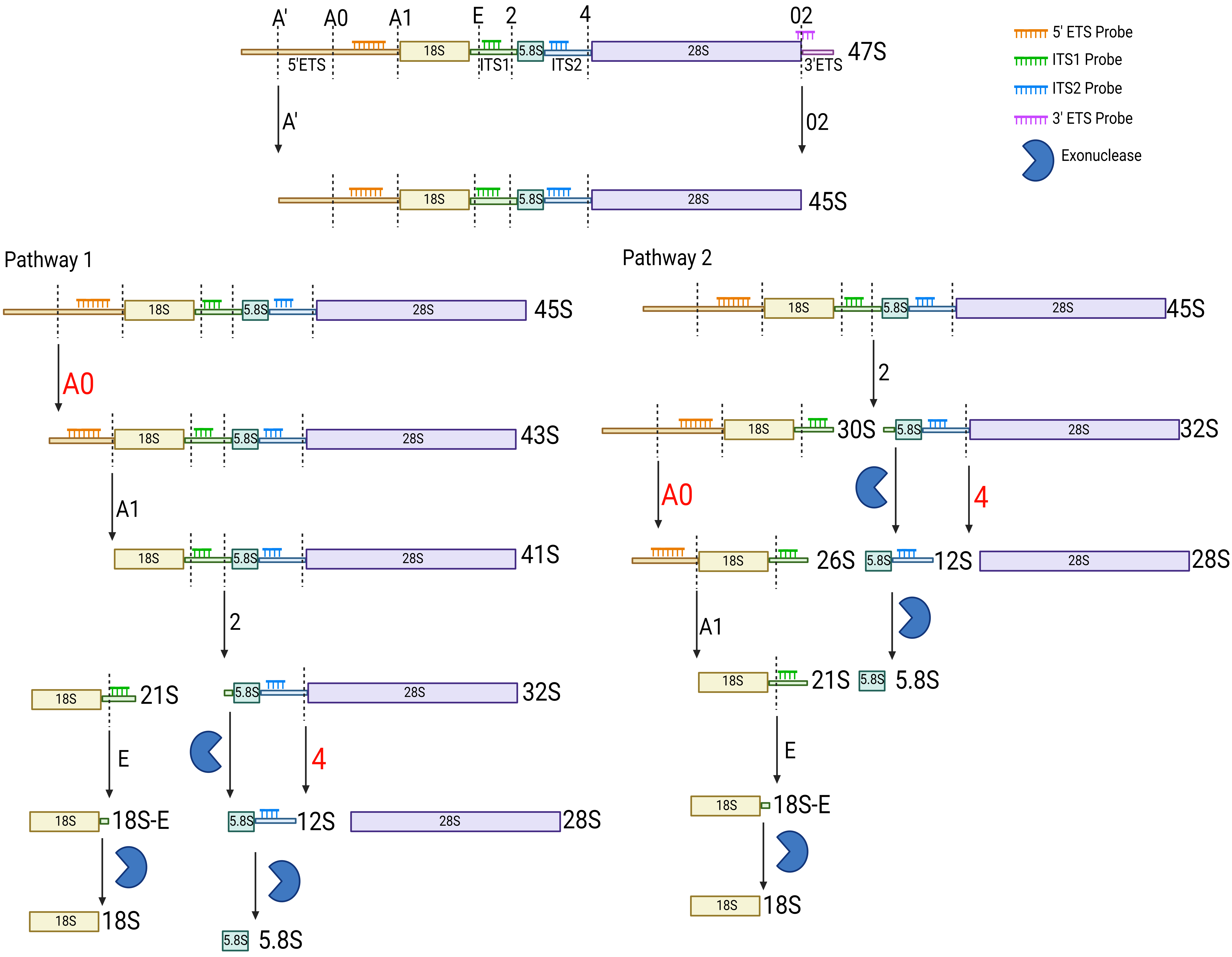
A'
A0
A1
E
2
4
02
18S
5.8S
28S
5'ETS
ITS1
ITS2
3'ETS
47S
45S
5' ETS Probe
ITS1 Probe
ITS2 Probe
3' ETS Probe
Exonuclease
Pathway 1
Pathway 2
43S
41S
21S
32S
30S
26S
12S
18S-E

**Supplementary Figure 3.** Overview of the pre-ribosomal RNA processing steps.

The diagram of pre-ribosomal RNA processing was based on the results of earlier studies, which have been previously summarized (3). The names of cleavage sites and their map location are shown within the map of the 47S pre-ribosomal RNA at the top of the figure. Map location of cleavage sites and probes is displayed throughout. Diagram made with www.biorender.com.

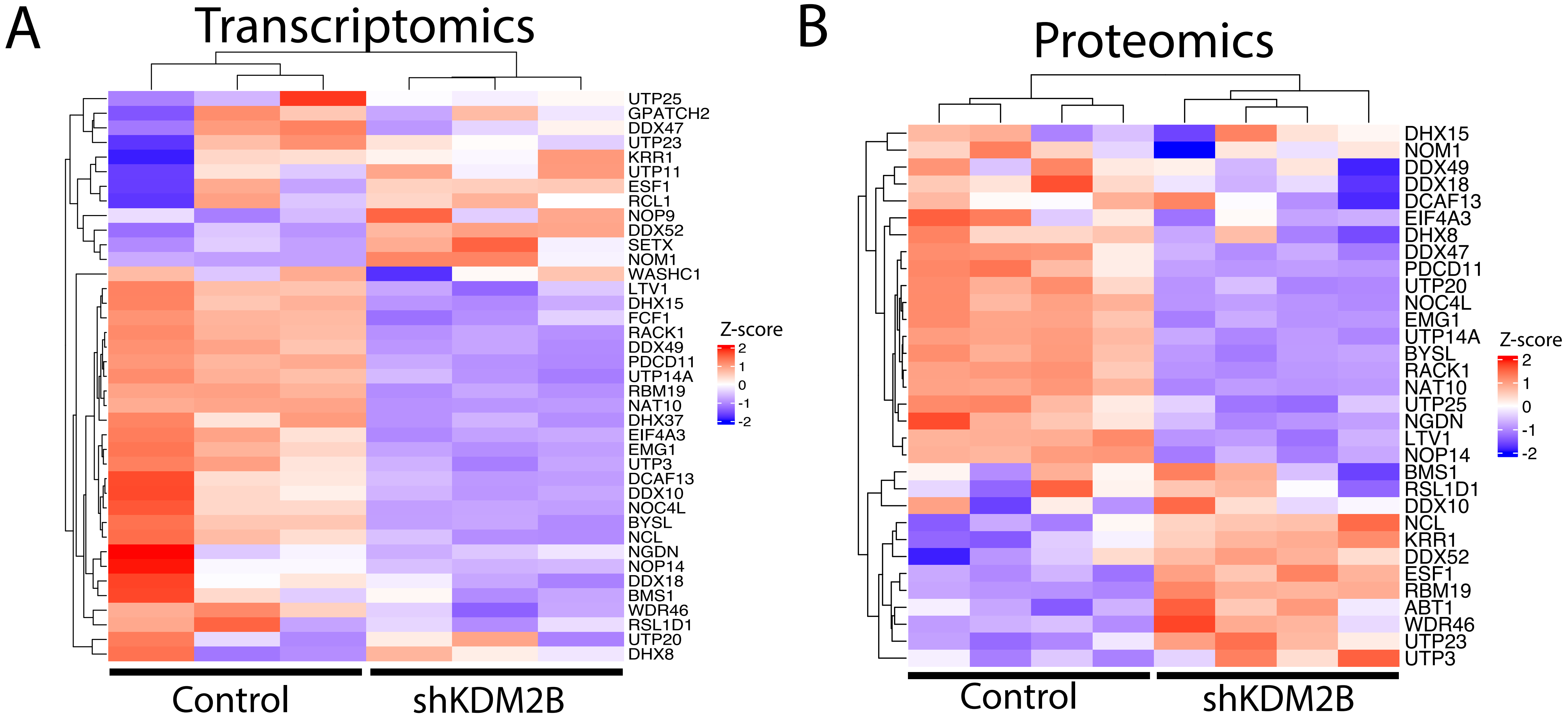
A
Transcriptomics
UTP25
GPATCH2
DDX47
UTP23
KRR1
UTP11
ESF1
RCL1
NOP9
DDX52
SETX
NOM1
WASHC1
LTV1
DHX15
FCF1
RACK1
DDX49
PDCD11
UTP14A
RBM19
NAT10
DHX37
EIF4A3
EMG1
UTP3
DCAF13
DDX10
NOC4L
BYSL
NCL
NGDN
NOP14
DDX18
BMS1
WDR46
RSL1D1
UTP20
DHX8
Z-score
2
1
0
-1
-2
Control
shKDM2B
B
Proteomics
DHX15
NOM1
DDX49
DDX18
DCAF13
EIF4A3
DHX8
DDX47
PDCD11
UTP20
NOC4L
EMG1
UTP14A
BYSL
RACK1
NAT10
UTP25
NGDN
LTV1
NOP14
BMS1
RSL1D1
DDX10
NCL
KRR1
DDX52
ESF1
RBM19
ABT1
WDR46
UTP23
UTP3
Z-score
2
1
0
-1
-2
Control
shKDM2B

**Supplementary Figure 4.** The knockdown of KDM2B impacts ribosome biogenesis factors involved in the assembly and maturation of the SSU processome.

The expression of genes encoding ribosome biogenesis factors involved in the assembly and maturation of the Small Subunit (40S) processome in control and shKDM2B MDA-MB-231 cells. **A.** Heatmap of RNA-seq data and **B.** Heatmap of TMT proteomics data. Heatmaps were made using the ComplexHeatmap R package (https://github.com/jokergoo/ComplexHeatmap), performing unbiased clusterings for the samples and genes. Signal intensities normalized for each gene, were based on Z-scores calculated form the geometric mean of each gene.

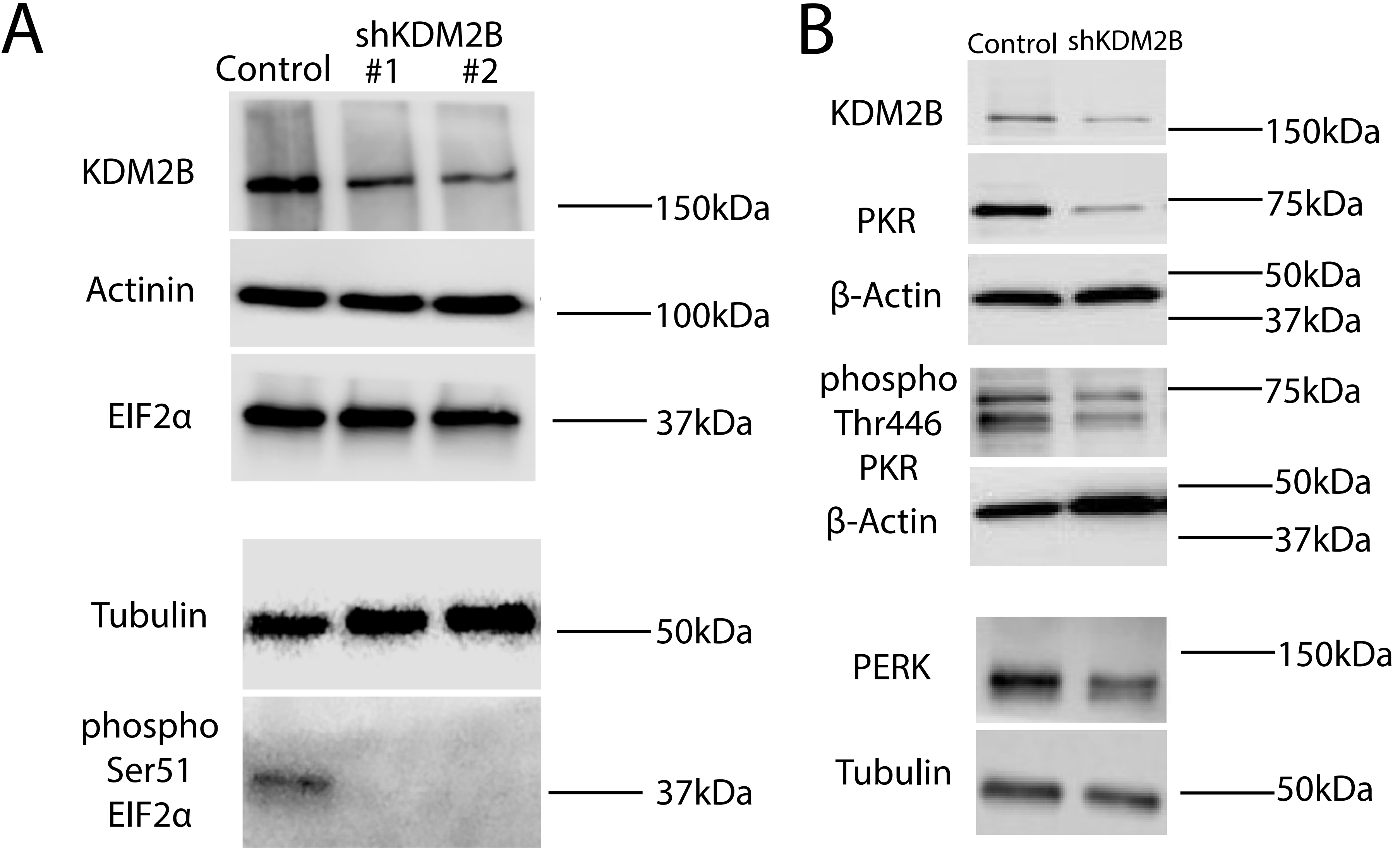

A
Control
shKDM2B
#1
#2
KDM2B
150kDa
Actinin
100kDa
EIF2α
37kDa
Tubulin
50kDa
phospho
Ser51
EIF2α
37kDa
B
Control
shKDM2B
KDM2B
150kDa
PKR
75kDa
β-Actin
50kDa
37kDa
phospho
Thr446
PKR
75kDa
β-Actin
50kDa
37kDa
PERK
150kDa
Tubulin
50kDa

**Supplementary Figure 5.** The knockdown of KDM2B relieves ISR and RQC in cancer cells expressing high levels of KDM2B.

Cell lysates of control or shKDM2B-transduced MDA-MB-231 cells were probed with **A.** KDM2B, EIF2α or phospho-Ser51 EIF2α specific antibodies and probed with **B.** KDM2B, PERK, PKR, or phospho-Thr446 PKR, specific antibodies.

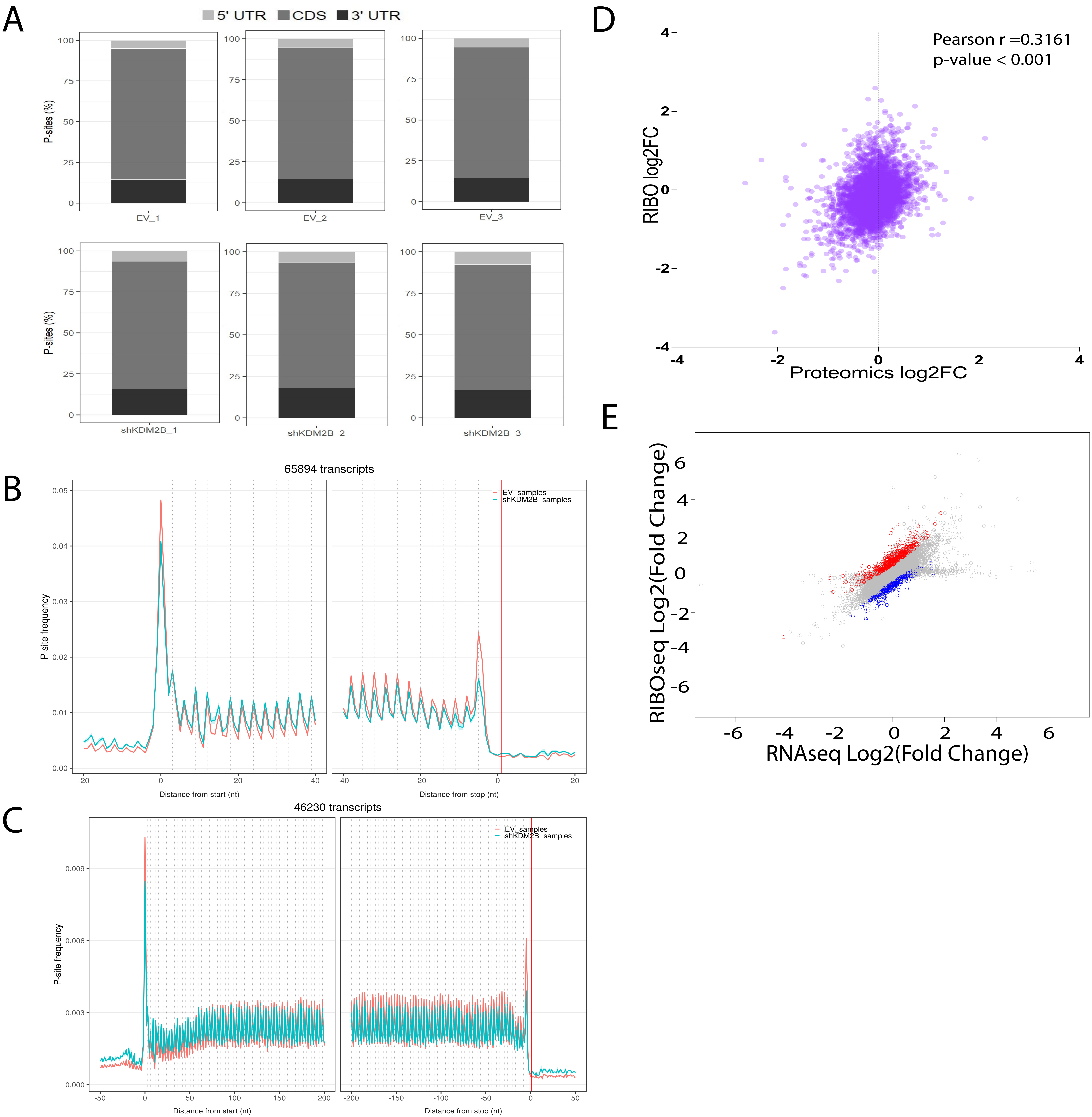
A
5' UTR
CDS
3' UTR
P-sites (%)
EV_1
EV_2
EV_3
shKDM2B_1
shKDM2B_2
shKDM2B_3
B
65894 transcripts
EV_samples
shKDM2B_samples
P-site frequency
Distance from start (nt)
Distance from stop (nt)
C
46230 transcripts
EV_samples
shKDM2B_samples
P-site frequency
Distance from start (nt)
Distance from stop (nt)
D
Pearson r =0.3161
p-value < 0.001
RIBO log2FC
Proteomics log2FC
E
RIBOseq Log2(Fold Change)
RNAseq Log2(Fold Change)

**Supplementary Figure 6.** P-site density of Ribo-seq across all samples

**A.** Distribution of P-site signals in the 5′ UTR, CDS, and 3′ UTR, genome-wide. **B. and C.** Periodicity of ribosome-protected fragments in control and shKDM2B-transduced MDA-MB-231 cells within 40 nt (~13 codons) or within 200 nt (~67 codons) from the translation start and stop codons. **D.** Correlation plot of Ribo-Seq and TMT proteomics data in shKDM2B-transduced MDA-MB-231 cells, for all the genes with data from both analyses. The x-axis shows the log2(fold change) in protein abundance, and the y-axis shows the log2(fold change) in the density of ribosome-protected RNA fragments in shKDM2B-transduced vs. control cells. Spearman correlation rho = 0.3161, p-value < 0.0001. **E.** Correlation plot of RNA-Seq and Ribo-Seq data in shKDM2B-transduced MDA-MB-231 cells. The x-axis shows the log2(fold change) in gene expression (reads per million), and the y-axis shows the log2(fold change) in the density of ribosome-protected RNA fragments in shKDM2B-transduced vs control cells. Dots representing individual transcripts were labeled in red if they exhibited significantly increased translational efficiency and blue if they exhibited significantly decreased translational efficiency in shKDM2B cells (Absolute log2 (foldchange) > 0.5 and adjusted p-value ≤ 0.05).

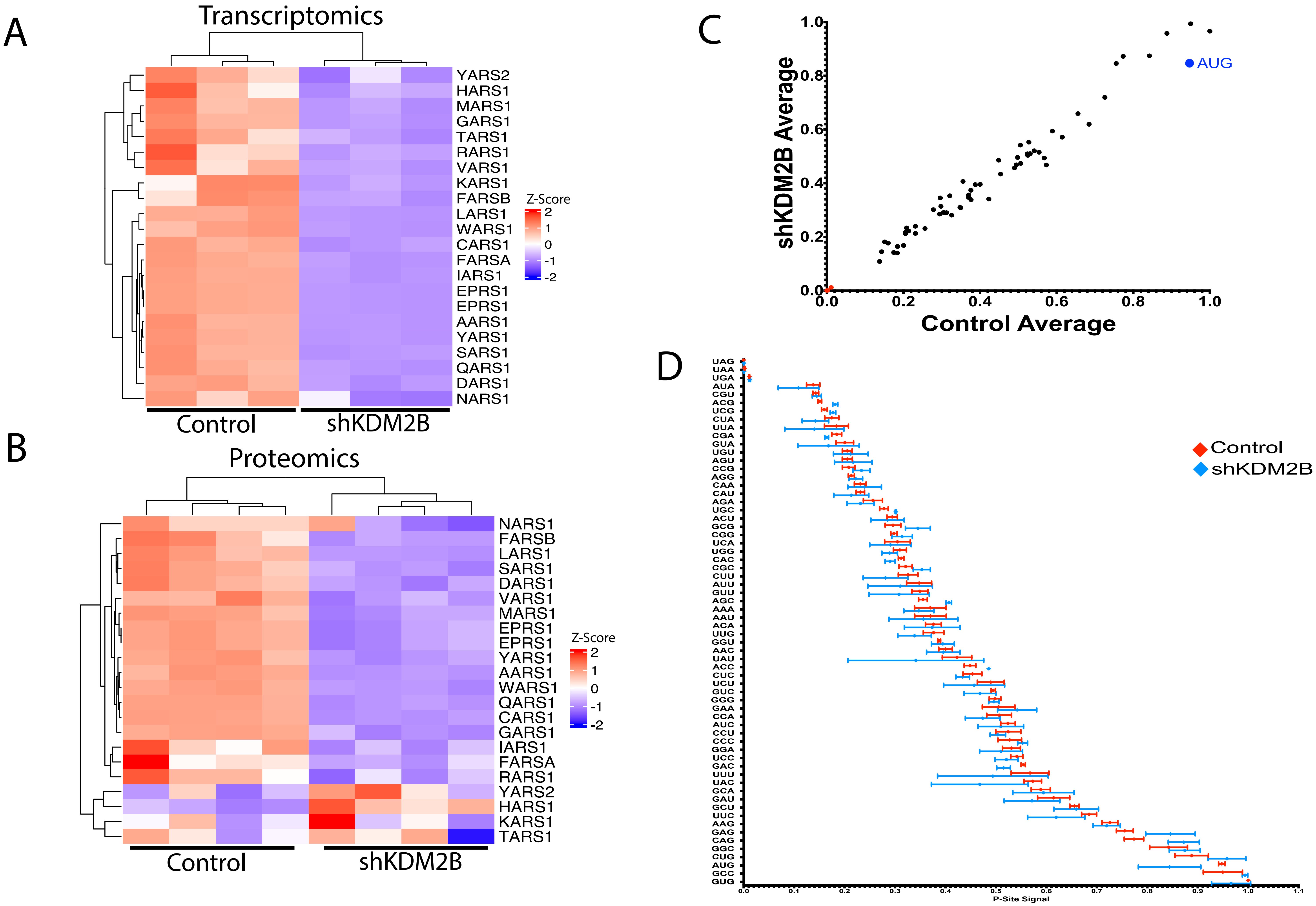

A
Transcriptomics
YARS2
HARS1
MARS1
GARS1
TARS1
RARS1
VARS1
KARS1
FARSB
LARS1
WARS1
CARS1
FARSA
IARS1
EPRS1
EPRS1
AARS1
YARS1
SARS1
QARS1
DARS1
NARS1
Z-Score
2
1
0
-1
-2
Control
shKDM2B
B
Proteomics
NARS1
FARSB
LARS1
SARS1
DARS1
VARS1
MARS1
EPRS1
EPRS1
YARS1
AARS1
WARS1
QARS1
CARS1
GARS1
IARS1
FARSA
RARS1
YARS2
HARS1
KARS1
TARS1
Z-Score
2
1
0
-1
-2
Control
shKDM2B
C
shKDM2B Average
Control Average
AUG
0.0
0.2
0.4
0.6
0.8
1.0
D
UAG
UAA
UGA
AUA
CGU
ACG
UCG
CUA
UUA
CGA
GUA
UGU
AGU
CCG
AGG
CAA
CAU
AGA
UGC
ACU
GCG
CGG
UCA
UGG
CAC
CGC
CUU
AUU
GUU
AGC
AAA
AAU
ACA
UUG
GGU
AAC
UAU
ACC
CUC
UCU
GUC
GGG
GAA
CCA
AUC
CCU
CCC
GGA
UCC
GAC
UUU
UAC
GCA
GAU
GCU
UUC
AAG
GAG
CAG
GGC
CUG
AUG
GCC
GUG
0.0
0.1
0.2
0.3
0.4
0.5
0.6
0.7
0.8
0.9
1.0
1.1
P-Site Signal
Control
shKDM2B

**Supplementary Figure 7.** Expression of amino acyltRNA synthetases and codon usage in KDM2B knockdown relative to control cells.

**A** and **B.** Heatmaps of the expression of Aminoacyl tRNA synthetase genes in control and shKDM2B-transduced MDA-MB-231 cells. **A.** RNA levels based on the analysis of RNA-Seq data, and **B.** Protein levels based on the analysis of TMT Proteomics data. Heatmaps were made using the ComplexHeatmap R package (https://github.com/jokergoo/ComplexHeatmap). Signal intensities were normalized based on Z-scores calculated from the geometric mean of each gene. **C.** Correlation plot of the average values of the P-site signal for each codon in control (x-axis) and shKDM2B (y-axis) cells. The start codon (AUG) is shown as a blue dot, and stop codons (UAA, UGA, and UAG) are shown as red dots. **D.** P-site signals (mean ± SD) for individual codons in control and shKDM2B-transduced MDA-MB-231 cells. Statistics were calculated, using the multiple unpaired t-test and correcting for multiple hypothesis testing with the Bonferroni-Dunn method.

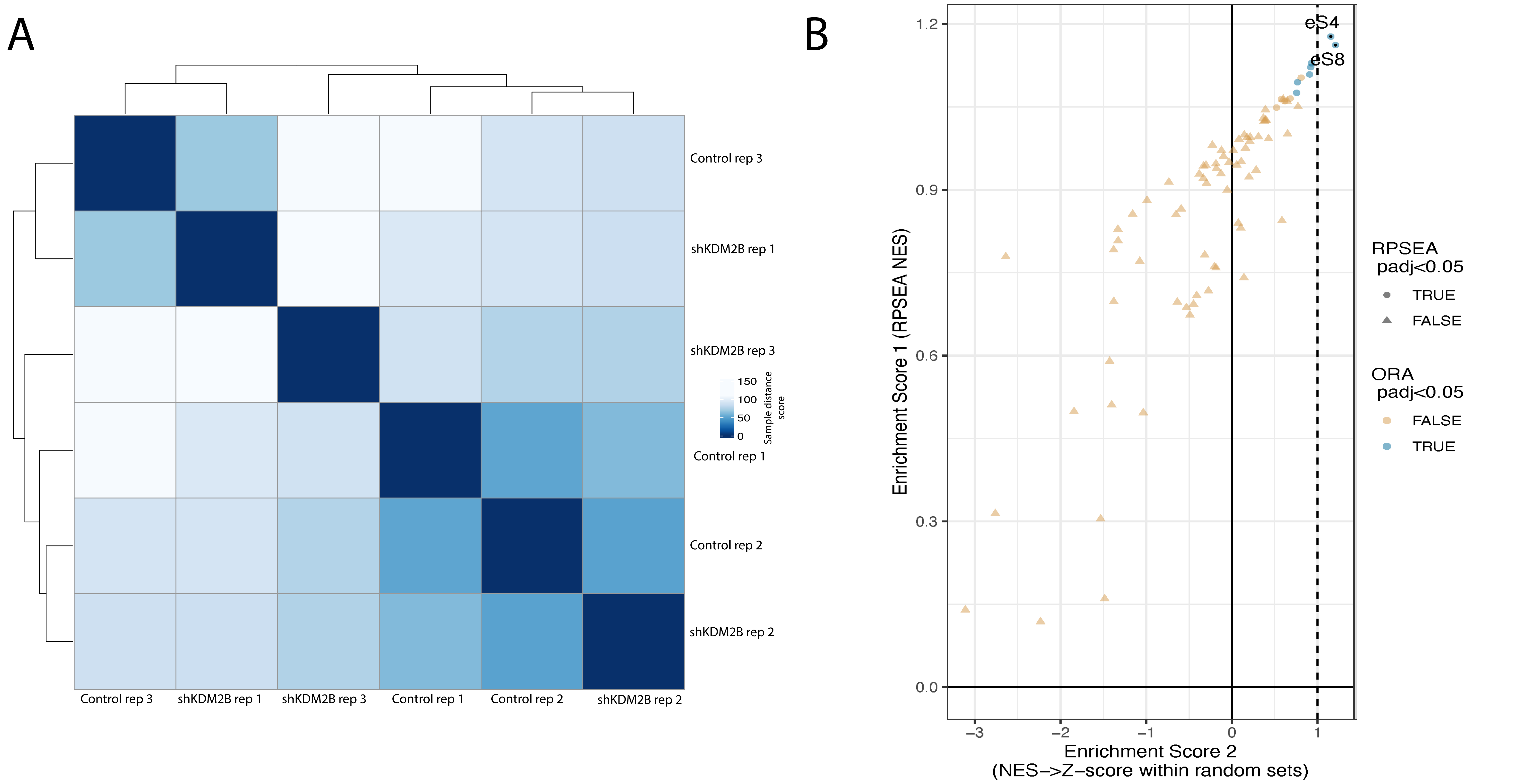
A
Control rep 3
shKDM2B rep 1
shKDM2B rep 3
Control rep 1
Control rep 2
shKDM2B rep 2
150
100
50
0
Sample distance score
B
eS4
eS8
1.2
0.9
0.6
0.3
0.0
−3
−2
−1
0
1
Enrichment Score 1 (RPSEA NES)
Enrichment Score 2
(NES−>Z−score within random sets)
RPSEA padj<0.05
TRUE
FALSE
ORA padj<0.05
FALSE
TRUE

**Supplementary Figure 8.** The knockdown of KDM2B does not have a robust ribosome heterogeneity signature.

DripARF ribosomal heterogeneity analysis (4). DripARF uses the abundance of Ribo-Seq-detected ribosomal RNA fragments to predict the abundance of ribosomal proteins that are known to bind these fragments in the ribosome. **A.** Matrix of hierarchically clustered individual samples. Clustering was based on the distance between samples, which was calculated by principal component analysis. **B.** Enrichment scores of rRNA fragments in shKDM2B relative to control cells, were calculated with two different methods and plotted.

A Translational Efficiency Downregulated de novo Motifs (STREME)

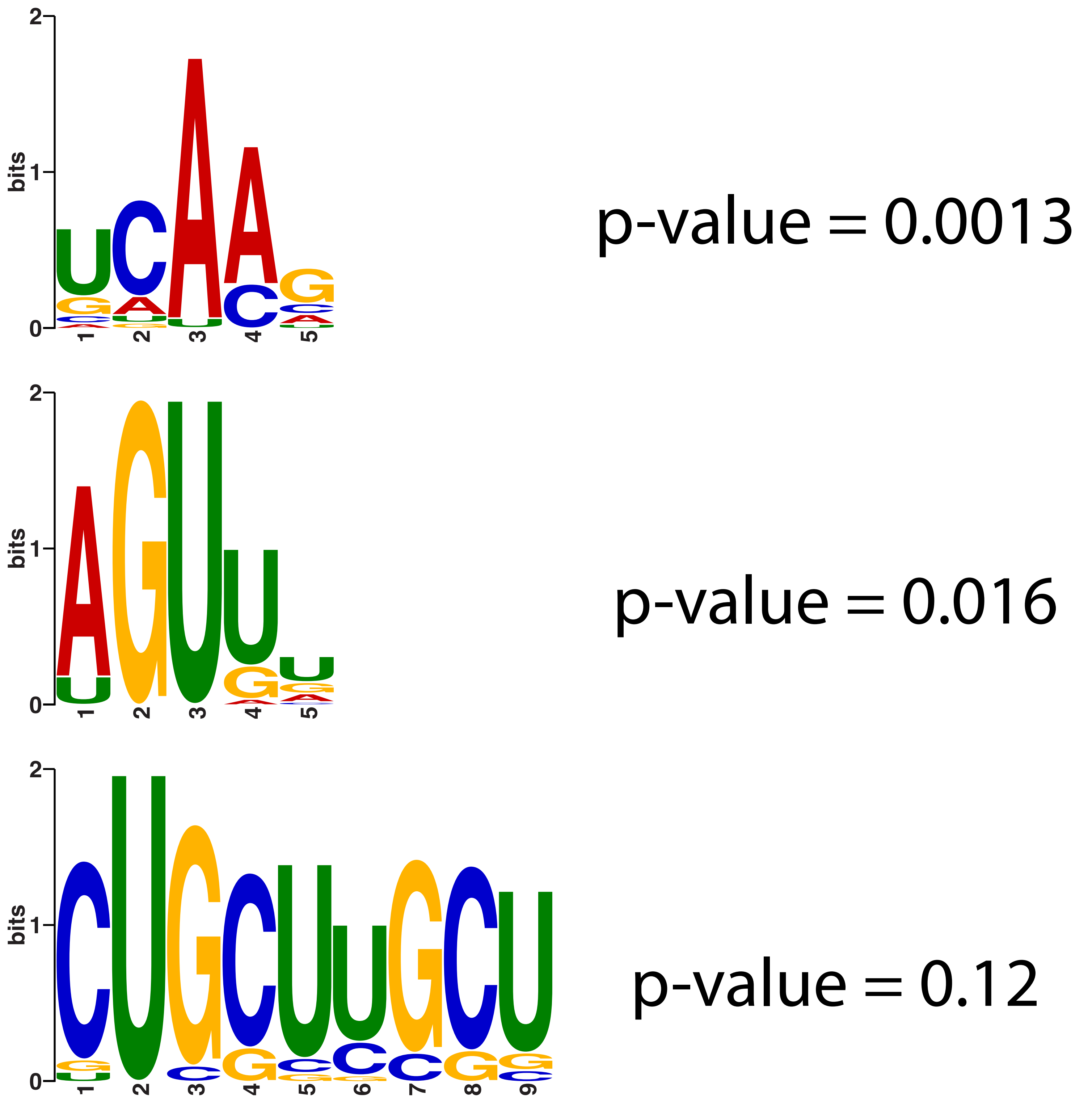


B Translational Efficiency Upregulated de novo Motifs (STREME)

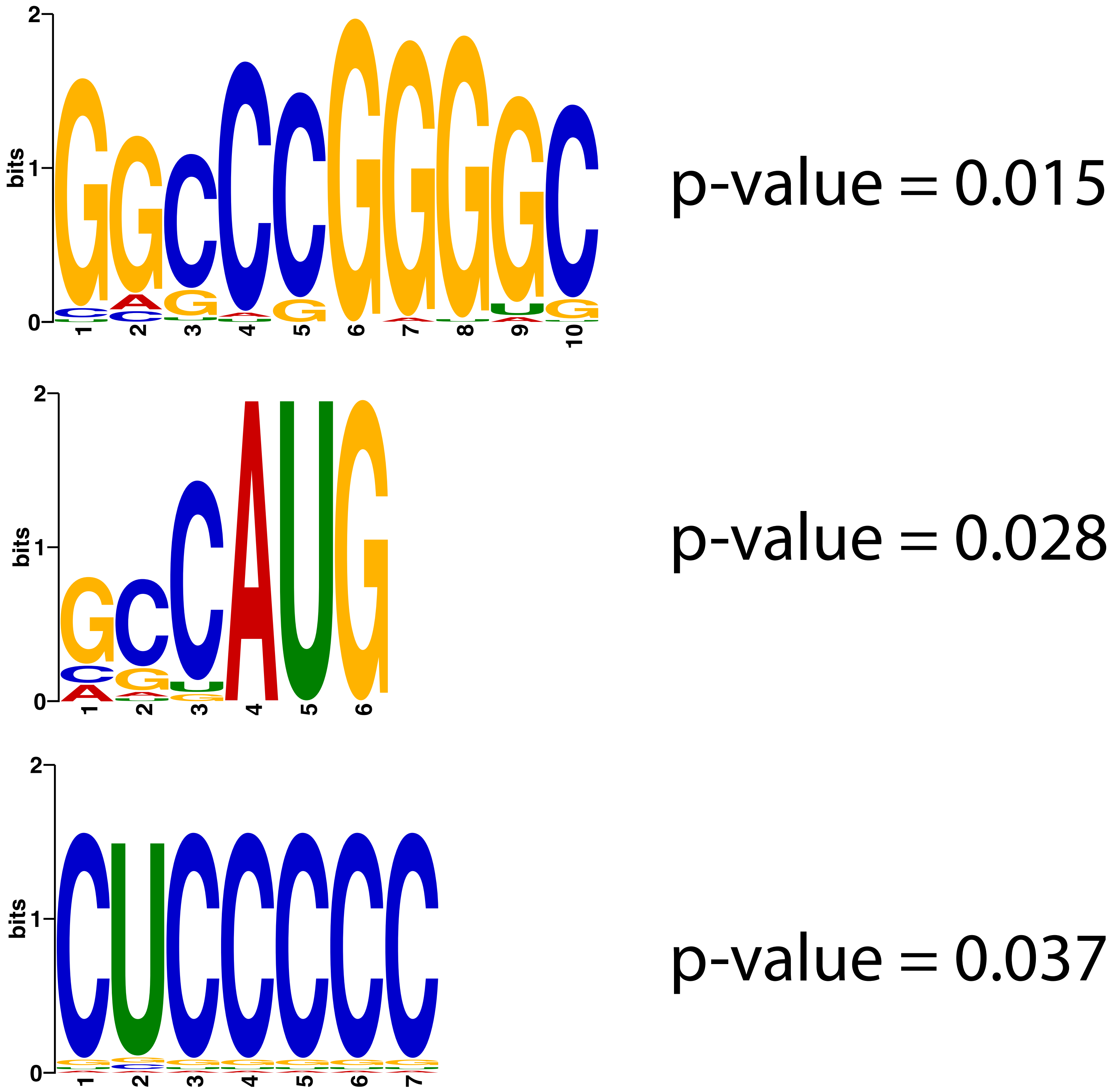


C Translational Efficiency Downregulated RNA Binding Motifs (AME)

| Logo | Database | ID | Alt ID | *p*-value |
|---|---|---|---|---|
| 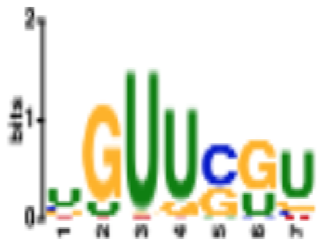 | Ray2013 rbp Homo sapiens | RNCMPT00164 | ZNF638 | 7.59e-7 |
| 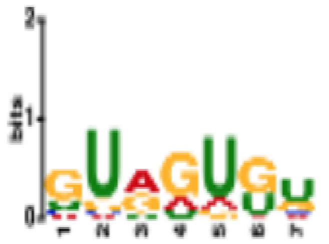 | Ray2013 rbp Homo sapiens | RNCMPT00177 | SFPQ | 3.30e-4 |
| 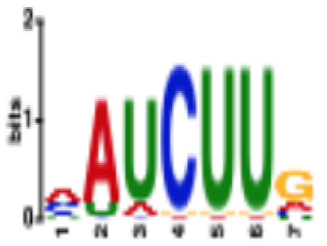 | Ray2013 rbp Homo sapiens | RNCMPT00037 | MATR3 | 5.20e-3 |
| 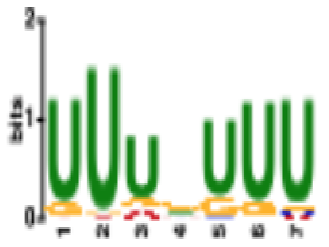 | Ray2013 rbp Homo sapiens | RNCMPT00086 | ZC3H14 | 9.91e-3 |
| 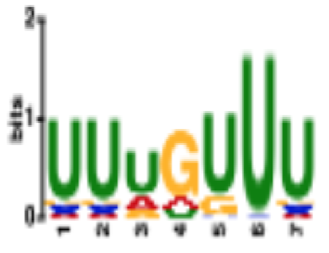 | Ray2013 rbp Homo sapiens | RNCMPT00112 | HuR | 1.15e-2 |

D Translational Efficiency Upregulated RNA Binding Motifs (AME)

| Logo | Database | ID | Alt ID | *p*-value |
|---|---|---|---|---|
| 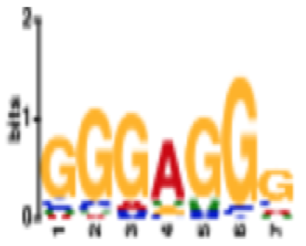 | Ray2013 rbp Homo sapiens | RNCMPT00160 | HNRNPH2 | 5.23e-11 |
| 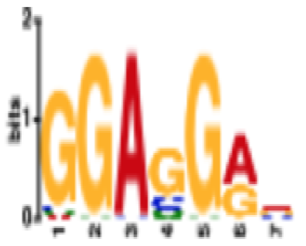 | Ray2013 rbp Homo sapiens | RNCMPT00109 | SRSF1 | 7.69e-9 |
| 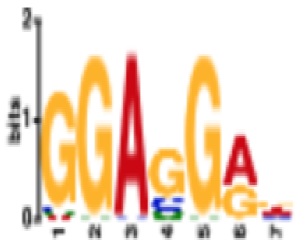 | Ray2013 rbp Homo sapiens | RNCMPT00108 | SRSF1 | 6.18e-8 |
| 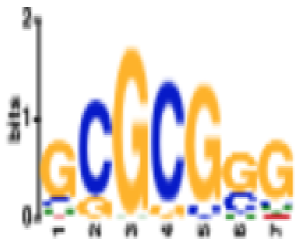 | Ray2013 rbp Homo sapiens | RNCMPT00113 | RBM4 | 1.37e-6 |
| 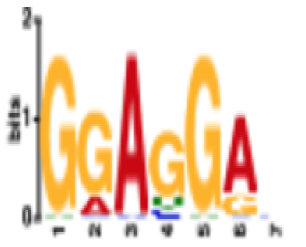 | Ray2013 rbp Homo sapiens | RNCMPT00107 | SRSF1 | 3.22e-6 |

**Supplementary Figure 9.** 5′ UTR Motif analysis of translationally regulated transcripts in control and shKDM2B-transduced cells.

**A** and **B**. De novo motif enrichment analysis (STREME) identified 5′ UTR sequence motifs enriched in transcripts that were translationally downregulated or translationally upregulated in shKDM2B-transduced MDA-MB-231 cells. **A.** Motifs enriched in translationally downregulated transcripts, and **B.** Motifs enriched in translationally upregulated transcripts. **C** and **D.** Motif enrichment analysis for known RNA binding protein motifs for **C.** Motifs enriched in translationally downregulated transcripts and **D.** Motifs enriched in translationally upregulated transcripts.

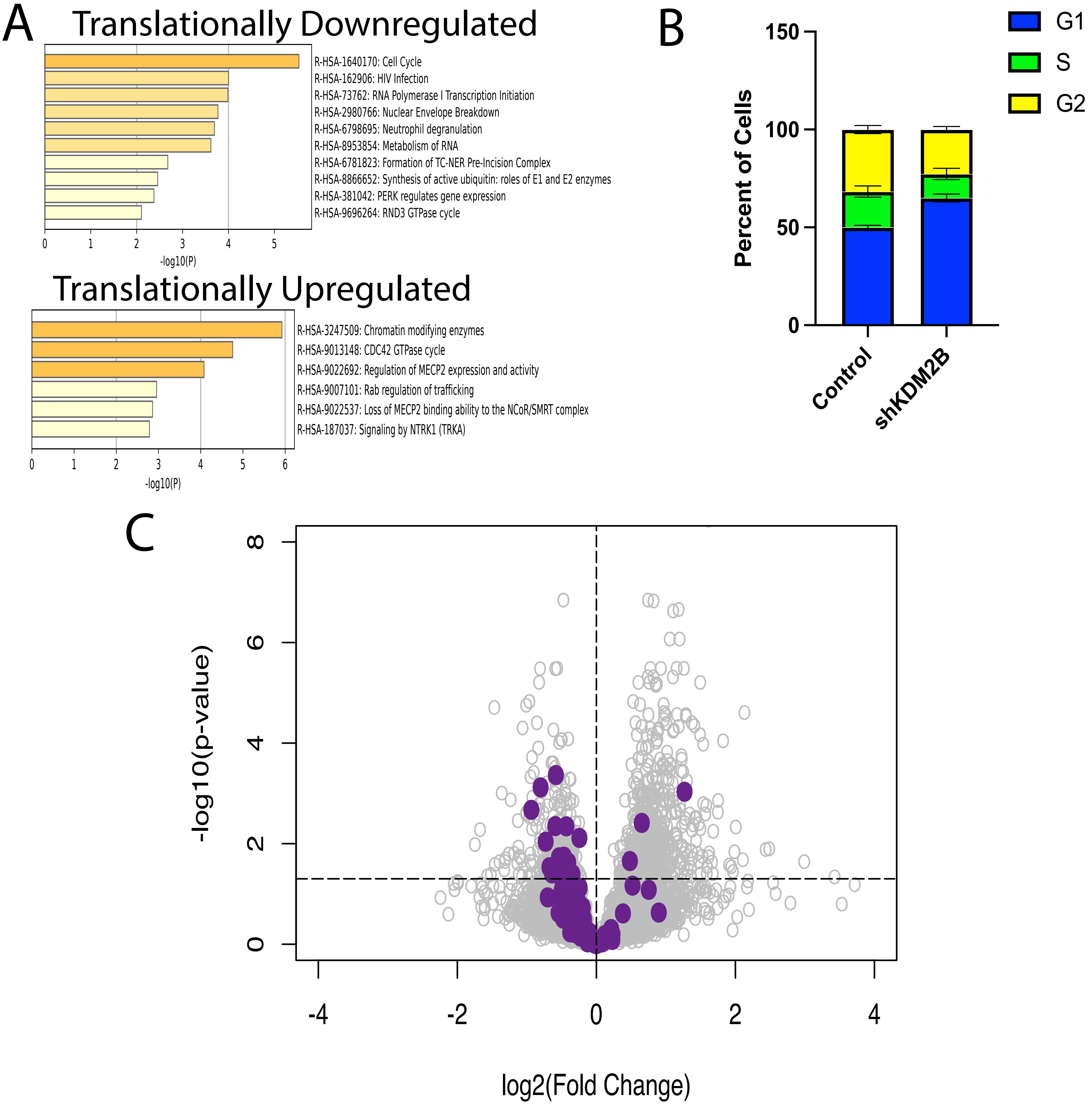
A
Translationally Downregulated
R-HSA-1640170: Cell Cycle
R-HSA-162906: HIV Infection
R-HSA-73762: RNA Polymerase I Transcription Initiation
R-HSA-2980766: Nuclear Envelope Breakdown
R-HSA-6798695: Neutrophil degranulation
R-HSA-8953854: Metabolism of RNA
R-HSA-6781823: Formation of TC-NER Pre-Incision Complex
R-HSA-8866652: Synthesis of active ubiquitin: roles of E1 and E2 enzymes
R-HSA-381042: PERK regulates gene expression
R-HSA-9696264: RND3 GTPase cycle
0 1 2 3 4 5
-log10(P)
Translationally Upregulated
R-HSA-3247509: Chromatin modifying enzymes
R-HSA-9013148: CDC42 GTPase cycle
R-HSA-9022692: Regulation of MECP2 expression and activity
R-HSA-9007101: Rab regulation of trafficking
R-HSA-9022537: Loss of MECP2 binding ability to the NCoR/SMRT complex
R-HSA-187037: Signaling by NTRK1 (TRKA)
0 1 2 3 4 5 6
-log10(P)
B
Percent of Cells
0 50 100 150
Control
shKDM2B
G1
S
G2
C
-log10(p-value)
0 2 4 6 8
-4 -2 0 2 4
log2(Fold Change)

**Supplementary Figure 10.** KDM2B controls the translational efficiency of mRNAs encoding regulators of the cell cycle.

**A.** Metascape enrichment analysis of transcripts that were Translationally downregulated (upper panel) or Translationally upregulated (lower panel) relative to all transcripts detected in the Ribo-seq study. **B.** Percentage (mean ± SD) of control and shKDM2B-transduced MDA-MB-231 cells in different phases of the cell cycle (G1, S, or G2/M) were determined by Propidium Iodide (PI) staining and flow-cytometry (3 biological replicates per condition). **C.** Volcano plot showing the log2(fold change) of the translational efficiency (ΔTE) of all Ribo-Seq-detected transcripts in shKDM2B-transduced relative to the control cells. The ΔTE log2(fold change) was plotted against the -log10 (p-value) of the (ΔTE) of all transcripts. The purple dots correspond to the ΔTE of transcripts, encoding proteins involved in the transition from G1 to S.
